# Deciphering the combinatorial expression pattern and genetic regulatory mechanisms of Beats and Sides in the olfactory circuits of *Drosophila*

**DOI:** 10.1101/2025.05.31.657193

**Authors:** Qichen Duan, Sumie Okuwa, Rachel Estrella, Chun Yeung, Yu-Chieh David Chen, Laura Quintana Rio, Khanh M. Vien, Pelin C. Volkan

## Abstract

Over the past decades, many critical molecular players have been uncovered to control distinct steps in olfactory circuit assembly in Drosophila. Among these, multi-member gene families of cell surface proteins are of interest because they can act as neuron-specific identification/recognition tags in combinations and contribute to circuit assembly in complex brains through their heterophilic or homophilic interactions. Recently, a multi-protein interactome has been described between the Beat and Side families of IgSF proteins. Here, we use the publicly available single-cell RNA-seq datasets and newly generated gene trap transgenic driver lines to probe the *in vivo* spatial expression pattern of the *beat/side* gene families in odorant receptor neurons (ORNs) and their synaptic target projection neurons (PNs). Our results revealed that each ORN and its synaptic target PN class expresses a class-specific combination of *beat/side* genes, hierarchically regulated by lineage-specific genetic programs. Though ORNs or PNs from closer lineages tend to possess more similar *beat/side* profiles, we also found many examples of divergence from this pattern among closely related ORNs and closely related PNs. To explore whether the class-specific combination of *beats/sides* defines ORN-PN matching specificity, we perturbed presynaptic *beat-IIa* and postsynaptic *side-IV* in two ORN-PN partners. However, disruption of Beat-IIa-Side-IV interaction did not produce any significant mistargeting in these two examined glomeruli. Though without affecting general glomerular targeting, knockdown of *side* in ORNs leads to the reduction of synaptic development. Interestingly, we found conserved expression patterns of *beat/side* orthologs across ORNs in ants and mosquitoes, indicating the shared regulatory strategies specifying the expression of these duplicated paralogs in insect evolution. Overall, this comprehensive analysis of expression patterns lays a foundation for in-depth functional investigations into how Beat/Side combinatorial expression contributes to the olfactory circuit assembly.

## Introduction

The *Drosophila* olfactory system provides an excellent model for understanding the genetic basis of the neuronal class-specific circuit organization and assembly. The first-order sensory neurons in the circuit are olfactory receptor neurons (ORNs), which are housed in the peripheral sensilla covering the antennae (Barish and Volkan, 2015; Brochtrup and Hummel, 2011; Hong and Luo, 2014; Jefferis and Hummel, 2006; Rodrigues and Hummel, 2008). Each ORN typically expresses only a single identity-defining olfactory receptor (OR) gene or a unique combination of up to three OR genes (Barish and Volkan, 2015). The cell bodies of ORNs expressing the same OR genes, thus of the same class, are dispersed across the antenna. Yet, their axons converge onto a single, uniquely positioned class-specific glomerulus in the antennal lobe, where they synapse with the second-order projection neurons (PNs) (Barish and Volkan, 2015; Hong and Luo, 2014). ORNs are thus defined by their OR expression and glomerular target identity, comprising ∼60 classes (Task et al., 2022). PNs, born from distinct neuroblast lineages surrounding antennal lobes, are also defined by their unique dendritic targeting of the dedicated glomeruli. At the same time, their axons extend to other higher brain regions, mushroom bodies, and lateral horns, which integrate the odor signal with other sensory cues and internal states to guide the animals’ behaviors (Hong and Luo, 2014; Jefferis et al., 2001; Li et al., 2018; Yu et al., 2010). Importantly, within the antennal lobe, ORNs and PNs form a one-to-one match in the class-specific glomerulus. This precisely controlled organization brings many interesting questions. How are these neurons born? What molecules mediate the communication between the neurons of the same class, different classes, or pre- and post-synaptic partners? How do these neurons acquire the necessary cell surface signaling to mediate these interactions that eventually ensure the formation of this stereotyped glomerular map? As this one-to-one structure maintains over evolution from insects to mammals (Imai et al., 2010), the easy and genetically tractable *Drosophila* is a great platform to determine the lineage-specific and combinatorial expression of cell surface proteins, their establishment, function, and evolution.

Many molecular players have been identified to participate in organizing the *Drosophila* olfactory circuits, ranging from the critical regulatory hubs at the top of the developmental hierarchy, like transcription factors instructing the lineage and/or the expression of a broad range of cell surface molecules (CSMs) (Komiyama et al., 2004; Komiyama et al., 2003; Li et al., 2020a; Tichy et al., 2008; Xie et al., 2022), to the executors at the bottom of the hierarchy, which are critical CSMs themselves. These include many members of large protein families, like leucine-rich repeat superfamily members Toll-6/Toll-7 (Ward et al., 2015), Fili (Li et al., 2025; Xie et al., 2019), and Capricious (Hong et al., 2009); Teneurins Ten-a and Ten-m (Hong et al., 2012; Mosca and Luo, 2014); Cadherin superfamily members Ncad (Hummel and Zipursky, 2004; Zhu and Luo, 2004), Flamingo (Arguello et al., 2021), and Fat2 (Kug) (Vien et al., 2023); Immunoglobulin Super Family (IgSF) members Dscam with extremely diverse isoform repertoire (Goyal et al., 2019; Hattori et al., 2007; Hummel et al., 2003; Zhu et al., 2006) and DIPs/Dprs, which consist of multiple paralogs (Barish et al., 2018). Our previous work showed that class-specific combinatorial expression of DIPs/Dprs organizes ORN axons within glomeruli (Barish et al., 2018). DIPs/Dprs are of particular interest because they form two multi-member subfamilies (11 DIPs and 21 Dprs), bind one another (Dprs to DIPs) primarily through heterophilic interactions, and exhibit striking cell-type-specific expression (Brovero et al., 2021; Carrillo et al., 2015; Cheng et al., 2019a; Cheng et al., 2019b; Cosmanescu et al., 2018; Özkan et al., 2013; Wang et al., 2022). Besides the olfactory system, several trans-synaptically interacting DIPs and Dprs have been shown to control synapse selectivity and formation in the visual circuits and neuromuscular junction (Ashley et al., 2019; Carrillo et al., 2015; Courgeon and Desplan, 2019; Dombrovski et al., 2025; Menon et al., 2019; Tan et al., 2015; Venkatasubramanian et al., 2019; Xu et al., 2019; Xu et al., 2022; Xu et al., 2018).

There are around 130 IgSF-encoding genes in the Drosophila genome (Sanes and Zipursky, 2020). In addition to well-characterized DIPs/Dprs comprised of 32 members, another two families, the Beaten path (Beat) family (14 paralogs) and Sidestep (Side) family (eight paralogs) proteins (Figure 1A), are less studied. They share many features with DIPs/Dprs. They also belong to IgSF, with two or five extracellular Ig domains mediating adhesion (Figure 1B), and form a heterophilic interaction network (Figure 1A) (Li et al., 2017b; Özkan et al., 2013). Beats/Sides have been shown to control the neuromuscular junction formation in both larval and adult motor systems (de Jong et al., 2005; Fambrough and Goodman, 1996; Heymann et al., 2022; Kinold et al., 2021; Kinold et al., 2018; Pipes et al., 2001; Siebert et al., 2009; Sink et al., 2001). Very recently, studies have begun to reveal their roles in synaptic specificity and induction to assemble the adult visual system (Carrier et al., 2025; Dombrovski et al., 2025; Osaka et al., 2024; Yoo et al., 2023). However, little is known about whether and how they contribute to the olfactory circuit organization. We observed that the expression levels of *beats/sides* in antennal tissues increase over development, and most of them tend to have higher transcriptional levels in the latter half of the pupal stage, from 40h after puparium formation (APF) throughout adulthood (Figure S2A), during which the stereotyped glomerular map is being formed. Given their protein properties and known roles in neural development, we sought to illustrate the *beats/sides*’ expression patterns and test their functions in building the *Drosophila* olfactory circuit.

**Figure 1.**
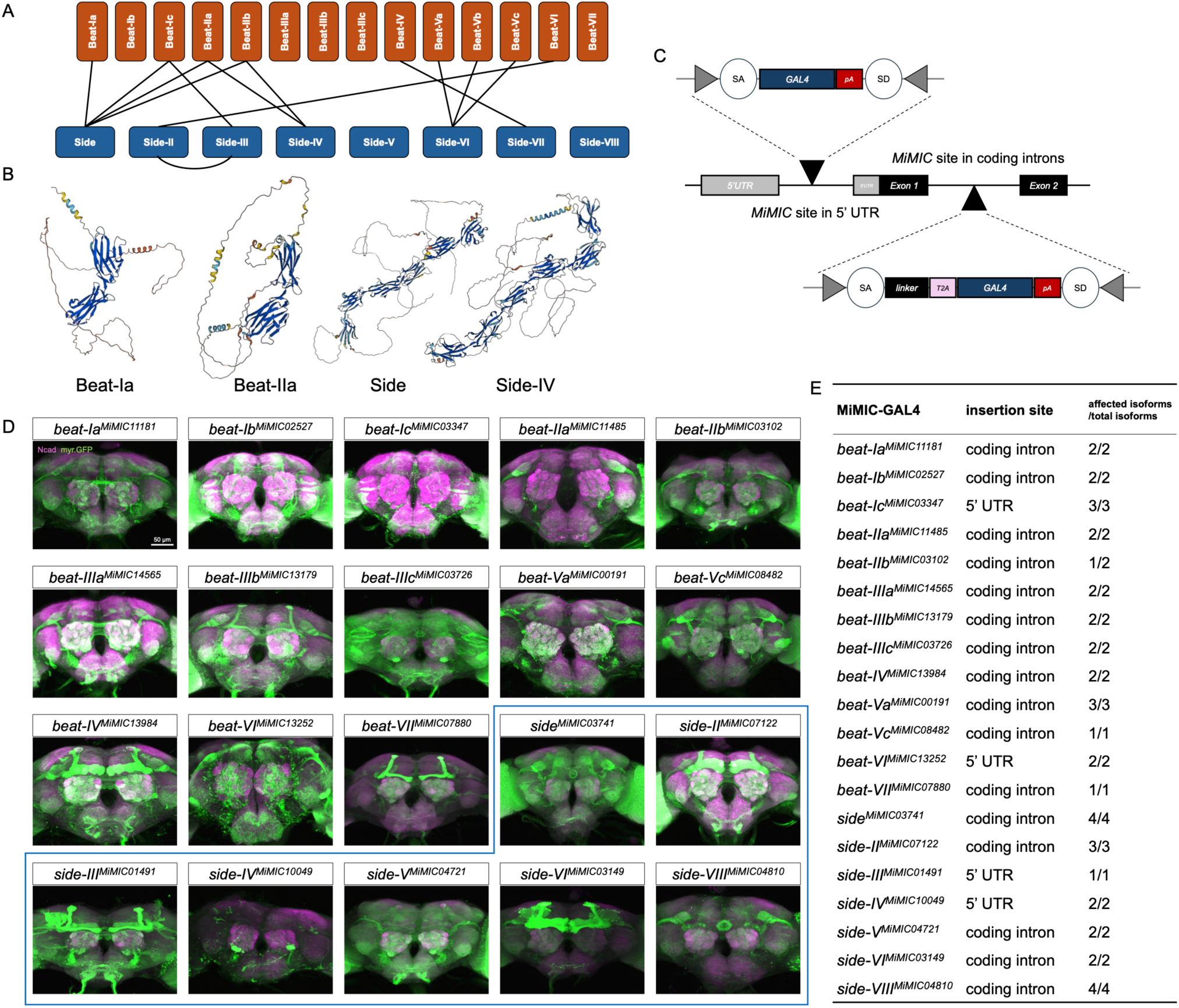
Beats/Sides are heterophilically interacting IgSF proteins. **(A)** Schematics showing the known interactions between Beat and Side proteins. **(B)** AlphaFold-predicted structures of example Beat and Side proteins. Beats have two extracellular Ig domains, while Sides’ extracellular parts consist of five Ig domains and one Fibronectin type III domain. Transmembrane and intracellular domains are poorly predicted. Structures were downloaded from AlphaFold.ebi.ac.uk (Jumper et al., 2021). **(C)** Schematic showing *MiMIC*-based insertion of the GAL4-coding construct into either the 5’ UTR or the “coding” intron, which will hijack the expression of the host gene. If *GAL4* is inserted into introns within the 5’ UTR, the *GAL4* artificial exon will first be kept from splicing and then will be translated as it has its own start codon. If the in-frame *T2A-GAL4* construct is inserted into an intron between two coding exons, this whole artificial exon will also be prevented from splicing and then translated along with the upstream coding sequence. During translation, the *T2A* sequence causes the ribosome to fail at synthesizing the peptide bond, and thereby, a truncated peptide of the host gene and a GAL4 protein will be produced separately. In both cases, GAL4 is expected to be expressed in the same cell where the native gene is expressed. SA, splice acceptor; SD, splice donor; pA, Hsp70 polyadenylation signal to terminate transcription; linker sequence ensures *T2A-GAL4* will be translated in frame. **(D)** Expression pattern of each *beat/side* in the brain revealed by *beat/side-MiMIC-GAL4*. Green is the GFP signal driven by the gene trap GAL4, and magenta is anti-Ncad staining showing the brain architecture. **(E)** Table summary of the insertion site and the isoforms labeled by each *beat/side-MiMIC-GAL4*.

In this study, we systematically characterized the expression pattern of *beat/side* family genes in both ORNs and PNs at single-cell and single-class levels. By analyzing the previously published single-cell RNA-seq datasets and genetically probing the native expression with *MiMIC*-based *beat/side* gene trap *GAL4* lines, we revealed that each ORN or PN class expresses a unique combination of *beats/sides*, and this *beat/side* profile is likely regulated by the lineage-specific genetic programs in a hierarchical manner. We also tested the functional relevance of one interacting pair, Beat-IIa and Side-IV, based on their matching expression pattern in partner ORNs and PNs. However, perturbation of either one pre- or post-synaptically did not result in apparent glomerular mistargeting. Nonetheless, we found pervasive ORN synapse defects when the *side* was knocked down in ORNs, suggesting its possible role in synaptic development. Interestingly, we found analogous expression principles for *beat/side* orthologs across ORNs in mosquitoes and ants, indicating the shared strategies across evolution for circuit assembly through lineage-specific regulation of cell surface protein combinations, particularly by coordinating the expression of multiple duplicated paralogs. Overall, our study reveals the combinatorial expression pattern and functional role that Beat/Side proteins play in olfactory circuit assembly and synaptic development.

## Results

### Genetically probing the *beat/side* expression *in vivo* by a collection of *MiMIC*-based gene trap driver lines

Recently, efforts to profile the transcriptional landscape of each cell across the whole fruit fly body have provided a valuable resource to examine the gene expression patterns in the cell types of interest. However, in the olfactory system, single-cell RNA-seq has only captured a limited portion of ORN or PN classes, leaving the transcriptome of many other ORN and PN classes unknown. To fully reveal the *beat/side* combinatorial profile, we used a *MiMIC*-based gene trap approach to generate a collection of transgenic *beat/side-*specific *GAL4* driver lines (Diao et al., 2015; Venken et al., 2011). By swapping the *GAL4* construct into 5’ UTR-located *MiMIC* sites or the in-frame *T2A-GAL4* construct into introns between two coding exons, we could make GAL4 hijack the expression of the host gene (Figure 1C and Materials and Methods). We then obtained *GAL4* driver lines for 13 of the 14 *beat* members and seven of the eight *side* members, except *beat-Vb* and *side-VII* (Figure 1D, E). Notably, *beat/side* genes generally have one to four annotated isoforms, and our *beat/side-MiMIC GAL4* collection is expected to trap all isoforms of each gene except *beat-IIb*, of which the *GAL4* only captures the expression of one of two isoforms (Figure 1E). This near-complete driver line collection reveals the remarkable enrichment of *beat/side* expression in neurons, from larval, pupal, to adult stages, in both peripheral and central nervous systems (Figure 1D; Figure S1). At the gross brain level, *beats/sides* are differentially expressed across different brain regions, including the antennal lobe, central complex, mushroom body, etc. (Figure 1D). This driver collection thus provides a valuable toolkit to study *beat/side* functions in diverse neuronal contexts in the olfactory circuits and beyond. Next, we used intersectional genetic strategies to restrict the reporter expression to ORNs or PNs, which allowed us to map the *beat/side* expression across glomeruli innervated by ORNs or PNs, respectively.

### A glomerular map of *beat/side* expression in ORNs

Our antennal bulk RNA-seq through pupal development shows that *beats/sides* are generally expressed at higher levels at later stages of glomerular formation, reaching the adult levels by mid-pupal stages (Figure S2A). To delineate the *beat/side* expression in each ORN class, we examined the publicly available single-ORN RNA-seq datasets from three distinct developmental stages: 24h APF (early pupal stage), 42-48h APF (mid-pupal stage), and adulthood (Li et al., 2020a; McLaughlin et al., 2021). At 24h APF, ORN axons have arrived at the antennal lobes and chosen a medial versus lateral antennal lobe trajectory, which positions them en route to their future glomerular regions. By the mid-pupal stage, the glomerular targeting is almost completed, and ORNs start forming synapses with their matching postsynaptic PNs. By the beginning of the adult stage, a stereotypical and discrete glomerular map has formed, and all glomeruli are now discernible. We visualized these datasets in bubble plots showing the fraction of positive cells and the mean expression of genes of interest of each ORN class (Figure S2B). This reveals that each ORN class possesses a unique signature of *beat/side* combinations throughout the developmental stages (Figure S2B). Combining bulk antennal RNA-seq and ORN single-cell RNA-seq results, we found that *sides* are generally expressed at higher levels in ORNs than *beats* (Figures S2A, B). Some *beats*, including *beat-Va*, *beat-Vb*, and *beat-Vc*, are barely expressed in any cell types within the antenna, whereas *side-VIII* appears to be predominantly expressed in non-neuronal cells of the antenna but not in ORNs (Figure S2A, B).

As many ORN classes are not captured in the single-cell RNA-seq datasets, we used the *MiMIC-GAL4* lines to probe the ORN expression of different *beats/sides in vivo* in 3-5-day-old adult brains as the proxy of their developmental expression. We leveraged the *eyeless*-driven FLP recombinase to excise the *STOP* cassette from *UAS-FRT-STOP-FRT-mCD8.GFP*, such that membrane-localized GFP expression by *beat/side-GAL4* can be restricted to ORNs. This way, we can label any glomeruli innervated by ORNs expressing each *beat/side* while excluding the signal from post-synaptic PNs, where *eyeless* has no expression. We generated a near-complete glomerular *beat/side* expression map across all ORN classes (Figure 2A). Consistent with bulk and single-cell RNA-seq, *beat-Ic*, *beat-Va*, and *side-VIII* are not expressed in any ORN classes. Some family members, like *beat-Ib*, *beat-VI*, and *side-IV,* are only sparsely expressed in very few ORN classes (Figure 2A). On the other hand, other *beats/sides* are expressed in a much broader pattern, though at varying levels among different ORN classes (Figure 2A).

**Figure 2.**
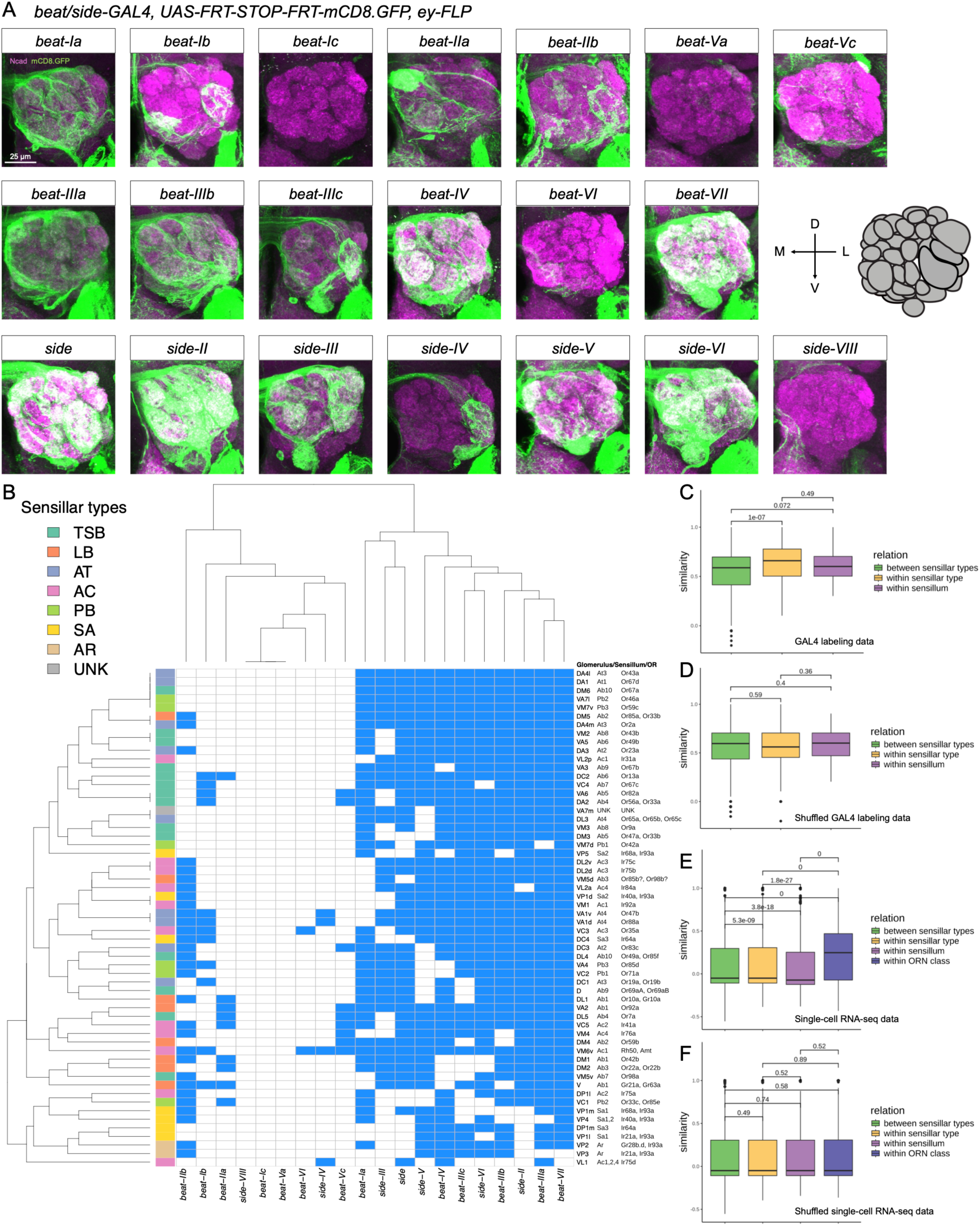
Lineage-dependent combinatorial expression of *beats/sides* across ORN classes. **(A)** A glomerular map of *beat/side* expression in ORNs revealed by transgenic *GAL4* driver lines. Glomerular structures are stained by anti-Ncad (magenta). Anti-GFP staining (green) highlights the glomeruli innervated by the indicated *beat/side*-positive ORNs. All antennal lobes shown are right antennal lobes. M, medial; L, lateral; D, dorsal; V, ventral. Each image is shown as the intensity projection over the Z-axis. **(B)** Hierarchical clustering of *beat/side* expression across ORNs based on the GAL4 labeling analysis. Row and column dendrograms are based on hierarchical clustering results. ORN class (named by its target glomerulus), the sensillum (sensillar subtype) where the ORN is housed, and the corresponding ORs expressed are shown on the right. The left row label is color-coded based on the ORN lineage. Blue means positive for the given gene expression, whereas white denotes negative. TSB, thin and small basiconics; LB, large basiconics; AT, antennal trichoids; AC, antennal coeloconics; PB, palp basiconics; SA, sacculus; AR, arista; UNK, unknown. VM6 glomerulus has been recently shown to be comprised of three subglomeruli, VM6v, VM6m, and VM6l, innervated by ORNs from Ac1 sensillum and Sacculus chamber III, respectively (Task et al., 2022). As it is difficult to distinguish Ac1-originated VM6v from the other two without additional colabeling, we counted VM6 as a single intact glomerulus for simplicity. **(C)** Similarity (Pearson’s correlation) of *beat/side* expression between ORN class pairs based on the expression matrix of (B) in three relation categories. **(D)** Similarity (Pearson’s correlation) of *beat/side* expression between ORN class pairs based on the expression matrix of (B), with row labels shuffled in three relation categories. **(E)** Similarity (Spearman’s correlation) of *beat/side* expression between ORN cell pairs based on the single-cell RNA-seq in four relation categories. **(F)** Similarity (Spearman’s correlation) of *beat/side* expression between ORN cell pairs based on the single-cell RNA-seq with ORN class annotation shuffled in four relation categories. In (C) to (F), P values from the Mann-Whitney U test of each comparison are shown.

Based on the GAL4 labeling patterns, we binarized the expression of all examined *beat* and *side* genes, as positive (value = 1) or negative (value = 0) for each gene in each ORN target glomerulus. We summarized the results and hierarchically clustered the genes by their expression pattern and ORN classes by their *beat/side-*expression profile, as shown in Figure 2B. We observed three types of gene expression patterns for *beat* and *side* genes across ORN classes: (1) broadly expressed, (2) expressed at a restricted pattern, or (3) not expressed (Figure 2B). More interestingly, ORN classes from the same lineage tend to be clustered based on the combinatorial *beat/side* expression (Figure 2B). This suggests a lineage-specific mechanism in patterning the ORN class-specific combinatorial expression of *beats/sides*. We decided to investigate this in further detail.

### Lineage-regulated genetic programs specify the combinatorial expression of *beats/sides* in ORNs

ORNs differentiate from precursor cells via a multi-step hierarchical genetic program (Barish and Volkan, 2015; Li et al., 2018). Early larval and pupal patterning factors first prepattern antennal imaginal discs (Royet and Finkelstein, 1997), followed by the sensillar type assignment by critical transcription factors, including Lozenge (Lz), Atonal, and Amos (Figure S2C) (Goulding et al., 2000; Gupta et al., 1998; Gupta and Rodrigues, 1997; zur Lage et al., 2003). We first examined the bulk antennal RNA-seq datasets profiling the transcriptional changes in mutants of *amos* (lacking trichoid and basiconic ORNs) or *atonal* (lacking coeloconic ORNs) (Menuz et al., 2014; Mohapatra and Menuz, 2019). We indeed found a couple of *beats/sides* that are downregulated in *atonal* and *amos* mutant antennae (Figure S2D), suggesting these *beats/sides* are enriched in the cells housed in Amos/Atonal-specified sensillar types. A few genes are also upregulated in *amos* and *atonal* mutants (Figure S2D). These *beat* and *side* genes are either transcriptionally upregulated or are now enriched in the mutants due to changes in ORN ratios.

Upon sensillar type selection, sensillar subtype fates are further specified by an additional set of transcription factors like Rotund (Rn), Dachshund (Dac), and Engrailed (En) (Figure S2C) (Barish and Volkan, 2015; Blagburn, 2008; Li et al., 2016; Li et al., 2013; Song et al., 2012). Finally, within each sensillar subtype (or sensillum), one multipotent precursor cell undergoes several sequential asymmetric divisions that eventually produce one to four different ORNs expressing distinct ORs and projecting to distinct glomeruli and four non-neuronal supporting cells (Barish and Volkan, 2015; Li et al., 2018). This step is achieved by iterative recruitment of Notch signaling bifurcating cell fates, coupled with other mechanisms like epigenetic modifiers and late transcription factors (Barish and Volkan, 2015; Chai et al., 2019; Endo et al., 2007) (Figure S2C). We here present a simplified decision tree to illustrate the kinships between ORNs and to compare *beat/side* combinations based on lineage relationships (Figure S2C): ORNs mapped to the same class, defined as “within ORN class”; ORNs of different classes but housed in the same sensillar subtype, described as “within sensillar subtype”; ORNs from different sensillar subtypes but belonging to the same sensillar type, namely, “within sensillar type”; and the furthest kinship, ORNs housed in distinct sensillar types, i.e., “between sensillar types”. We compared the similarity of *beat/side* combinatorial expression between cells falling into these ORN kinship categories to obtain insights into the developmental regulation of *beat/side* expression.

Firstly, based on our gene-ORN class expression matrix derived from the genetic labeling data (Figure 2B), we calculated the similarity (Pearson’s correlation) of the *beat/side* expression vector between each ORN class pair, and found significantly higher similarity between ORN classes belonging to the same sensillar types than those from different sensillar types (Figure 2C). In contrast, this pattern is not observed from the randomized control, where we shuffled the ORN class identity with the expression vector (Figure 2D). Additionally, we found that this difference also holds for the single-cell RNA-seq datasets. By calculating the similarity (measured by Spearman’s correlation) according to gene expression between each single ORN, we found that, as expected, ORNs mapped to the same class display the highest pairwise similarity of *beat/side* profile in all stages (Figure 2E; Figure S2E). Moreover, ORNs within the same sensillar type appear to possess more similar *beat/side* profiles than ORNs from different sensillar types, in contrast to the shuffled control (Figure 2E, F), supporting that the lineage-intrinsic mechanisms set the *beat/side* expression.

However, we found that *beat/side* combinatorial expression tends to diverge between ORNs housed in the same sensillar subtype (Figure S2E). We observed even lower pairwise similarity according to the *beat/side* expression between these cells than between cells from different sensillar subtypes but sharing the identical sensillar type (Figure S2E). This observation stands true for all three stages (Figure S2E). In contrast, at 24h APF and mid-pupal stage, the similarity of the pan-CSM profile and the whole transcriptome linearly increases with closer kinship (Figure S2E). It suggests that the overall transcriptional profile and the cell surface gene expression patterns are primarily set by lineage-specific factors, such that cells with closer kinship tend to have more similar transcriptomic profiles and overall cell surface codes. Furthermore, *beats/sides* expression adds new levels of complexity to cell surface signals, potentially diversifying ORN-specific glomerular decisions, particularly among the most developmentally related ORNs in the same sensillar subtype. Interestingly, at the adult stage, this divergence also exists for overall CSM and cell-specific transcriptional profiles (Figure S2E). These results suggest that at the end of development, ORNs with the closest kinship may have acquired broader transcriptional variance, which may support the establishment of discrete glomerular maps necessary for olfactory transduction, odor discrimination, and processing to drive odor-guided behaviors.

While the single-cell RNA-seq-based analysis only reflects the trend for the limited cell types captured in that dataset, our genetic labeling-based comprehensive analysis of *beat/side* across all ORN classes reveals the existence of both convergence and divergence: ORNs from the same sensillar subtype exhibit binary *beat/side* similarity: they could be clustered together, suggesting their shared *beat/side* combinatorial expression; they could also be segregated, indicating the divergence of *beat/side* profile from genetically very close ORNs (Figure 2B). We therefore propose a model: in some sensillar subtypes, at the step of ORN terminal selection, *beat/side* combinatorial expression is diversified from a putative initial lineage-specific “template” to ensure the closely related ORNs gain more variation in these cell surface molecular repertoire, whereas in some sensillar subtypes, *beat/side* profile is pre-specified and these closely related ORNs show similar *beat/side* combinations.

Collectively, our thorough characterization of *beat/side* expression from both single-cell RNA-seq and gene trap *GAL4* intersectional labeling suggests that Beats/Sides could be the cell surface “executors” underlying the hierarchical genetic programs that specify the ORN fate, likely participating in fine-tuning the ORN wiring.

### A glomerular map of *beat/side* expression in PNs

ORNs synapse with dedicated PNs within each glomerulus to assemble the olfactory circuit, and Beats/Sides are involved in synaptogenesis (Osaka et al., 2024; Yoo et al., 2023). We also need to survey the *beat/side* expression on the PN side, which may inform whether Beats/Sides could function trans-synaptically to mediate ORN-PN matching. We started with the bulk PN RNA-seq (Li et al., 2020b) from the FACS-sorted PNs at 36h APF and adult stage. In general, *beats/sides* appear to be expressed at commensurate levels between these two time points (Figure S3A). Next, we mined the published single-cell RNA-seq data (Li et al., 2017b; Xie et al., 2021) for developing PNs to gain insights into the PN type-specific expression of *beats/sides*. This dataset captured PNs at 0h APF, 24h APF, 48h APF (mid-pupal), and the adult stage. Some PNs are born embryonically and are part of the larval olfactory circuit (Jefferis et al., 2001; Lin et al., 2012; Marin et al., 2005; Yu et al., 2010). At the beginning of the pupal stage, these PNs first prune their terminal axons and dendrites and re-extend their neurites to be integrated into the adult olfactory circuit following other larvally born PNs (Marin et al., 2005). From 0h to 24h APF, PNs project their axons to the mushroom body and lateral horn and their dendrites to the antennal lobes. PNs’ dendrites create a prototypical glomerular map before ORN axons arrive in the antennal lobes at around 24h APF. Thereafter, PNs begin to match their presynaptic ORN partners. And starting from the mid-pupal stage, they build synapses, refine the discrete glomerular organization, and finally mature the olfactory circuit by the end of the pupal stage (Jefferis et al., 2001). In terms of PN class-specific expression of *beats/side*s at different time windows, we found: (1) *beats/sides* are enriched at 24h APF and thereafter, while expressed at minor levels at the beginning of metamorphosis (Figure S3B); (2) compared with ORNs, *beats/sides* are generally expressed in a broader pattern, and all *beats/sides* are expressed in PNs while some *beats* and *side-VIII* are undetectable in ORNs (Figures S2B, S3B). Suppose Beats/Side mediate adhesion between ORNs and PNs; this observation raises a hypothetical model where broadly and less specifically located postsynaptic surface “locks” are matched by sparsely and more specifically distributed presynaptic surface “keys”.

Next, we used these *GAL4* lines to drive the GFP reporter expression in postsynaptic PNs by removing the *STOP* cassette from *UAS-FRT-STOP-FRT-mCD8.GFP* with the PN-specific *GH146-flippase*. We found generally broader expression of *beats/sides* across PN-labeled glomeruli, with a few sparsely distributed ones, like *beat-Ic* and *side-IV* (Figure 3A). We then summarized the binary expression of all *beats/sides* examined across all glomeruli identified (Figure 3B). Hierarchical clustering of PN types according to their *beat/side* expression shows, in general, a correlation between *beat/side* combination and their PN lineages (Figure 3B). We therefore further investigated this.

**Figure 3.**
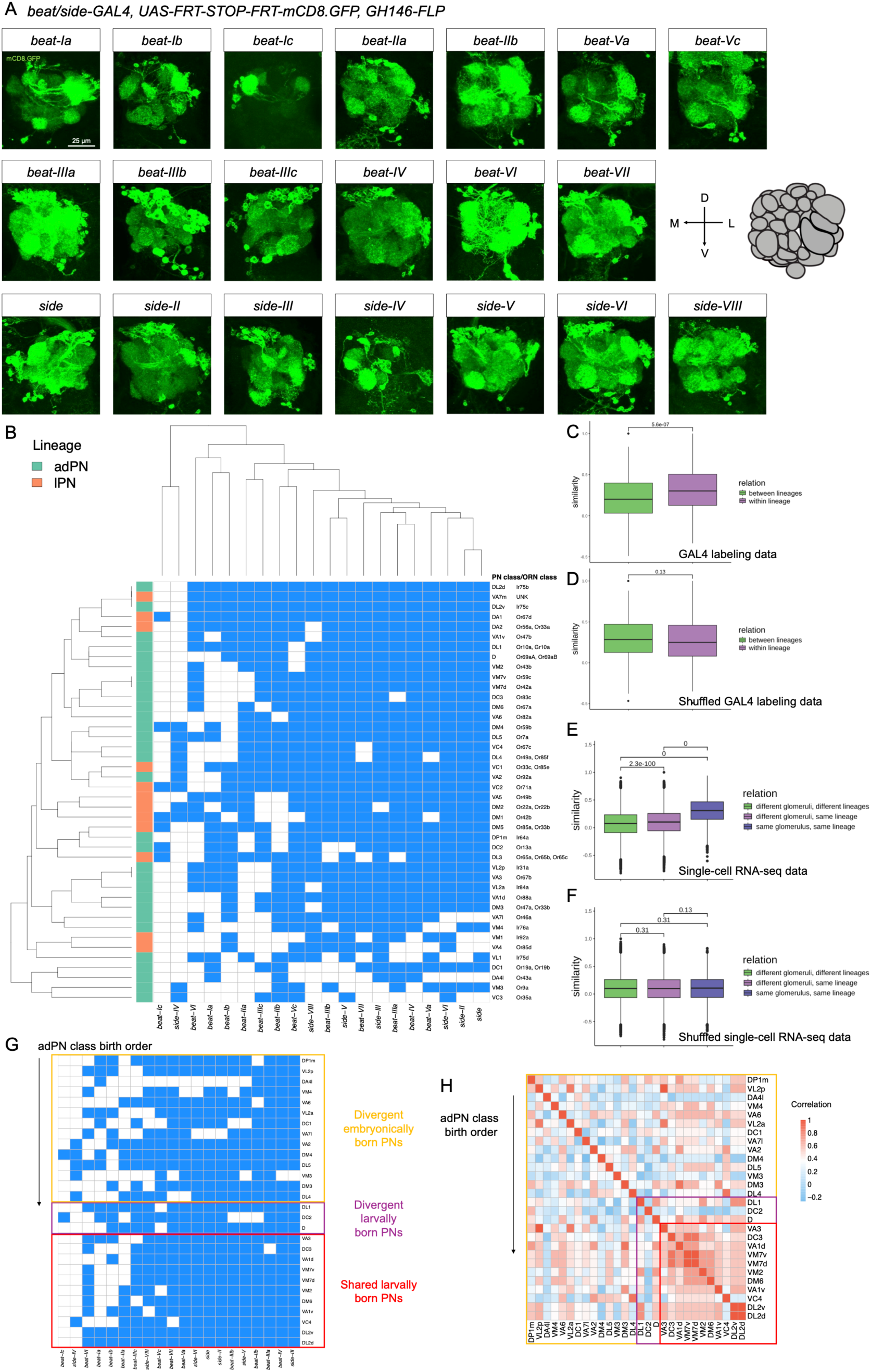
Lineage-correlated combinatorial expression of *beats/sides* across PN classes. **(A)** A glomerular map of *beat/side* expression in PNs revealed by transgenic *GAL4* driver lines. Only the GFP signal (green) highlights the glomeruli innervated by the indicated *beat/side*-positive PNs. Neuropils were stained with anti-Ncad antibody to help determine the glomerular identity but are not shown for visualization contrast. All antennal lobes shown are right antennal lobes. M, medial; L, lateral; D, dorsal; V, ventral. Each image is shown as the intensity projection over the Z-axis. **(B)** Hierarchical clustering of *beat/side* expression across PNs based on the GAL4 labeling analysis. Row and column dendrograms are based on hierarchical clustering results. PN class and its matching ORN class (named by corresponding ORs expressed) are shown on the right. The left row label is color-coded based on the uniglomerular PN lineage. Blue means positive for the given gene expression, whereas white denotes negative. **(C)** Similarity (Pearson’s correlation) of *beat/side* expression between PN class pairs based on the expression matrix of (B) in two relation categories. **(D)** Similarity (Pearson’s correlation) of *beat/side* expression between PN class pairs based on the expression matrix of (B) with row labels shuffled in two relation categories. **(E)** Similarity (Spearman’s correlation) of *beat/side* expression between PN cell pairs based on the single-cell RNA-seq in three relation categories. **(F)** Similarity (Spearman’s correlation) of *beat/side* expression between PN cell pairs based on the single-cell RNA-seq with ORN class annotation shuffled in three relation categories. In (C) to (F), P values from the Mann-Whitney U test of each comparison are shown. **(G)** The summary matrix is replotted from (B), but only PN classes from the adPN lineage with decoded birth order are shown. Rows representing PN classes are ordered based on the known birth sequence, while *beat/side* genes are hierarchically clustered. **(H)** Heatmap showing the Pearson’s correlation matrix between PN classes based on the *beat/side* expression. PN classes are the same as (G) and are ordered based on the birth sequence.

### Lineage-regulated genetic programs specify the combinatorial expression of *beats/sides* in PNs

PN fates are specified separately from ORNs by a sequential genetic program from three distinct neuroblast lineages (Figure S3C) (Jefferis et al., 2001; Li et al., 2018). A common neuroblast sequentially produces one ganglion mother cell and a self-renewed neuroblast. The ganglion mother cell further divides into one terminal PN, which innervates a dedicated glomerulus, and another daughter cell, which undergoes cell death (Lin et al., 2012; Lin et al., 2010). The younger neuroblast continues this process. Thus, different PN classes are born in a sequentially stereotyped order (Jefferis et al., 2001; Lin et al., 2012; Yu et al., 2010). Based on this, we defined the relationship between PNs into three categories: (1) PNs from the same lineage targeting the same glomerulus, (2) PNs from the same lineage targeting different glomeruli, and (3) PNs from different lineages targeting different glomeruli.

We again asked whether the *beat/side* profile similarity between PN classes is based on their developmental kinship. Calculating the PN class-specific *beat/side* expression similarity based on the *GAL4* labeling expression matrix showed higher similarity between PN classes of the same lineage than of different lineages (Figure 3C, D). Similarity analysis for *beat/side* expression in the single-PN RNA-seq datasets also confirmed these observations (Figure 3E, F; Figure S3D). In the same lineage, as expected, PNs targeting the same glomeruli also have more similar *beat/side* expression than PNs targeting different glomeruli (Figure S3D). This is also true for the expression of CSMs and the cellular transcriptional profiles at all four stages (Figure S3D). This suggests that, as part of their cell surface repertoire, the *beat/side* profiles in PNs are primarily set by the lineage and PN class-specific genetic mechanisms.

As PNs in each lineage are born in a stereotyped order, we also examined whether PN birth order determines its *beat/side* profile. We focused on the adPN lineage and calculated the correlation between each PN pair. We plotted their *beat/side* profile and sorted the PN classes in this lineage based on the reported birth order (Figure 3G). We also plotted the Pearson’s correlation coefficient between each pair of PN classes (Figure 3H). We found that the early-born PNs have more divergent *beat/side* profiles while later-born PNs gradually obtain a shared Beat/Side combinatorial “template” which gets less complex as new PN fates are generated (Figure 3G, H). This trend indicates that PN precursor-specific factors set the *beat/side* expression potential of daughter PNs and undergo a temporal fate specification along with the birth sequence of PNs. It is also interesting that, similar to ORNs (Figure 2B, C), closely related PNs can possess shared or divergent *beat/side* combinatorial expression, suggesting that different intrinsic genetic programs may be present in different precursor cells to control the opposing *beat/side* expression similarity in their daughter cells.

### Beats/Sides form a robust molecular interaction network between ORN and PN partners

The first identified Beat-Ia and Side pair act trans-synaptically to promote synapse formation at the neuromuscular junction (Fambrough and Goodman, 1996; Kinold et al., 2021; Kinold et al., 2018; Siebert et al., 2009; Sink et al., 2001). Recent work also identified additional Beat-Side interactions, including Side-IV and Beat-IIa/b, mediating neuronal recognition, connection specificity, and synaptogenesis in the fly visual system (Osaka et al., 2024; Yoo et al., 2023). We, therefore, hypothesized that Beat and Side interactions between ORNs and PNs can also regulate the synaptic matching. Guided by the known Beat-Side interactions based on the *in vitro* biochemical assays, we examined the *beat/side* expression between partner ORN and PN targeting the same glomerulus (Figures 1A, 4A). This analysis revealed a putative Beat/Side-mediated interactome between ORNs and PNs, which is promiscuous, redundant, and complex.

While several putative Beat-Side interactions exist between partner ORN and PN classes, we found one elegant and straightforward case: presynaptic Beat-IIa and postsynaptic Side-IV. *beat-IIa* is expressed in just a few ORN classes, including Or42b ORNs targeting one posterior glomerulus DM1 and Or92a ORNs targeting one anterior glomerulus VA2 (Figure 4B). In parallel, *side-IV* expression is restricted in PNs, including PNs projecting to DM1 and VA2 glomeruli (Figure 4C). Next, we tested whether perturbing *beat-IIa* expression in ORNs or *side-IV* expression in PNs results in glomerular targeting defects. To do this, we used a pan-ORN driver, *peb-GAL4,* to knock down *beat-IIa* in all ORNs while labeling Or42b ORNs or Or92a ORNs as a readout of glomerular targeting. However, we did not observe any gross phenotype in glomerular integrity, morphology, position, or ectopic mistargeting (Figure 4D, E). We also used a PN-specific driver, *GH146-GAL4,* to express *side-IV RNAi* and test if Or42b or Or92a ORNs target their glomeruli normally. Again, we found no visible differences between knockdowns and controls (Figure 4F, G). These results suggest that either Beat-IIa/Side-IV interaction in matching ORNs and PNs is unnecessary for glomerular targeting or is compensated by other Beat and Side proteins with redundant functions expressed in the same neurons.

**Figure 4.**
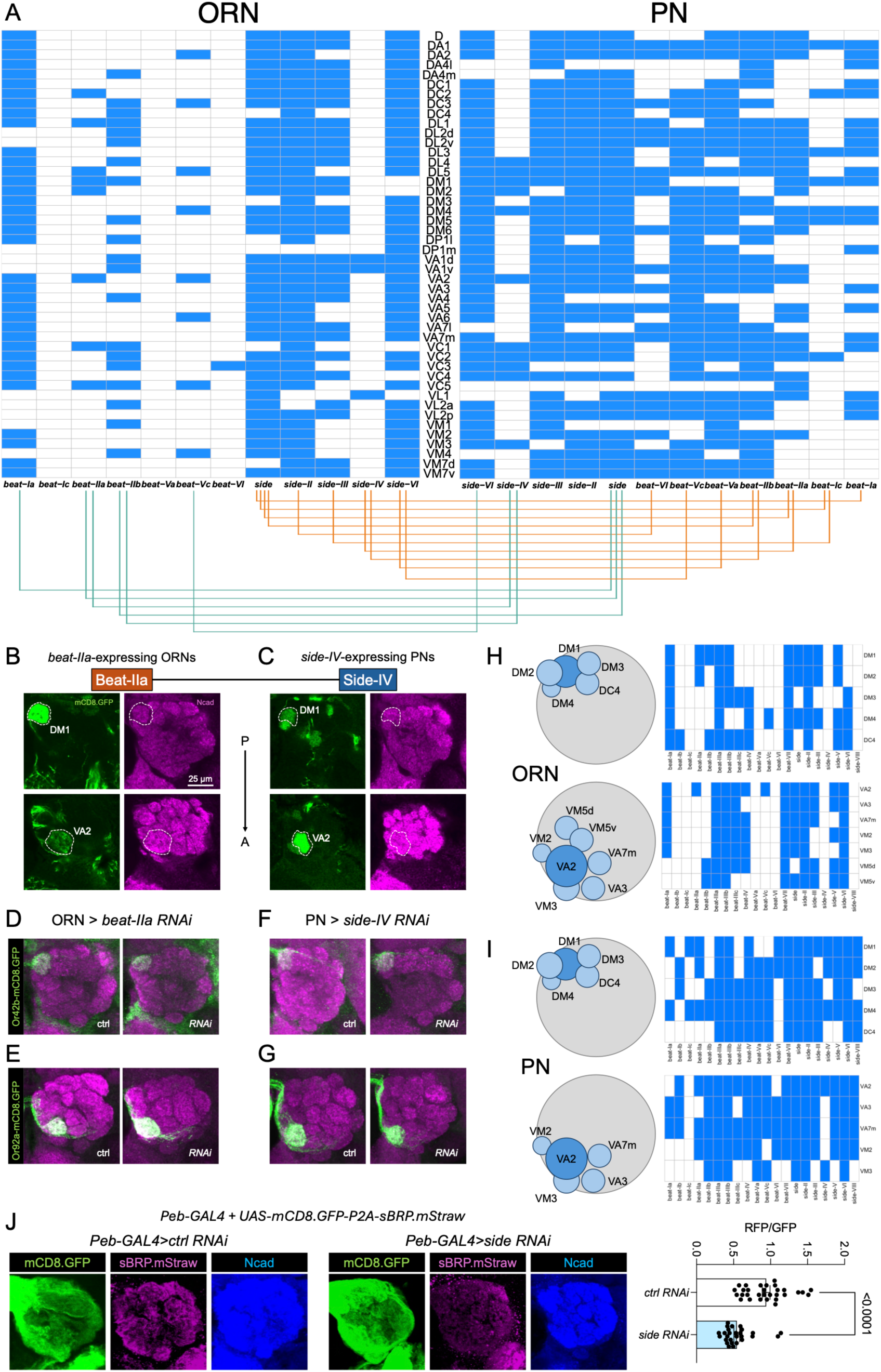
Beats-Sides interactions between matching ORN-PN partners can form a redundant, error-tolerant surface “barcode”, enabling recognition robustness and mediating synaptogenesis. **(A)** Potential trans-synaptic Beat/Side interactions between matching ORN-PN classes targeting each glomerulus. The ORN (left) and PN (right) profile of beat/side expression is based on Figures 2B and 3B, but it is replotted to highlight the ORN class and PN class innervating the same glomerulus. Based on *in vitro* biochemical characterization (Figure 1A), known binding interactions between Beats and Sides are indicated as lines. Green lines represent Beats -> Sides interactions, and red lines represent Sides -> Beats interactions (ORN -> PN). **(B)** ORNs targeting DM1 and VA2 glomeruli expressing *beat-IIa*, labeled by the intersection of *beat-IIa-T2A-GAL4* and *ey-FLP* to drive *UAS>STOP>mCD8.GFP*. Scale bar representing 25 µm also applies to each antennal lobe panel in (C) to (G) and (J). **(C)** PNs targeting DM1 and VA2 glomeruli expressing *side-IV*, labeled by the intersection of *side-IV-GAL4* and *GH146-FLP* to drive *UAS>STOP>mCD8.GFP*. In (B) and (C), the left columns show anti-GFP staining (green) and the right columns show anti-Ncad staining (magenta). **(D, E)** Knockdown of *beat-IIa* in ORNs by a pan-ORN driver, *peb-GAL4,* while labeling Or42b ORNs targeting DM1 glomerulus (D) and Or92a ORNs targeting VA2 glomerulus (E). **(F, G)** Knockdown of *side-IV* in PNs by a PN driver *GH146-GAL4* while labeling Or42b ORNs targeting DM1 glomerulus (F) and Or92a ORNs targeting VA2 glomerulus (G). Neuropils were stained with anti-Ncad antibody, shown in magenta. Representative images from 7 to 12 brains examined in each condition are shown (D-G). **(H)** The Beat/Side “barcode” in ORNs targeting DM1 or VA2 and their neighboring glomeruli, based on the *beat/side-GAL4* labeling results in Figure 2B. **(I)** The Beat/Side “barcode” in PNs targeting DM1 or VA2 and their neighboring glomeruli based on the *beat/side-GAL4* labeling results in Figure 3B. VM5d/v PN expression is unknown because these glomeruli are not labeled by the PN-FLP we used. In (H) and (I), the schematics show the antennal lobe (gray circle) and the relative positions of the indicated glomeruli (blue circles). **(J)** Representative antennal lobes and quantification showing the effects of knockdown of *side* on gross glomerular synaptic density, visualized by the membrane marker GFP and presynaptic marker RFP. Each dot is a single antennal lobe. N = 30 and 27 antennal lobes, respectively. P-value from the unpaired t-test between groups is shown.

Indeed, the combinatorial *beat/side* codes can distinguish ORNs targeting DM1 or VA2 glomeruli from ORNs innervating the neighboring glomeruli (Figure 4H), such that removing *beat-IIa* in ORNs doesn’t ambiguate the CSM repertoire and still preserves a level of diversity among the local axon fibers in a glomerular neighborhood. Similarly, PNs targeting DM1 or VA2 also harness this robust combinatorial “barcode”, and deleting *side-IV* in PNs may not abolish the dendritic diversity in the glomerular neighborhood (Figure 4I). To test for general defects in glomerular organization, we also conducted a pan-ORN driver *peb-GAL4* driven *UAS-RNAi* screening against *beat* and *side* genes that are expressed in ORNs (Figure S4A). This screen revealed mild defects in the VA1v glomerulus targeted by Or47b ORNs by knockdown of *beat-IIb, beat-IIIa/b/c,* and *side-III*, and a low-penetrance phenotype of split DM3 glomerulus targeted by Or47a ORNs (Figure S4A, B). However, additional controls showed that the VA1v phenotypes of *beat-IIIa/b/c* perturbation appeared to arise from the transgenic background that sensitizes the VA1v glomerular disorganization (Duan et al., 2023)(Figure S4C). These negative results might be because the Beat/Side combinatorial expression can theoretically act as a robust cell surface recognition code for each ORN or PN class, given its error-tolerance feature.

Given the lack of glomerular targeting defects, we next asked whether *beat* and *side* genes have roles in regulating synapse development and maintenance within glomeruli. For this, we used a previously generated transgenic reporter, *UAS-SynLight*, which expresses the presynaptic active zone marker, Bruchpilot-Short (sBRP), and the neurite membrane marker mCD8.GFP in the same transcript (Aimino et al., 2023). We expressed this transgene in ORNs to label and quantify the synaptic density in the antennal lobes while doing perturbation in ORNs. We calculated the ratio of the mean fluorescence intensity of RFP and GFP in the antennal lobe as a proxy of synaptic density (Figure S5A). This way normalizes the potential variability in transcription levels, imaging conditions, or neuron size across groups. As inhibiting neural activity has been reported to decrease synapse numbers, we first silenced ORN neuronal activity by overexpressing the mutant *shaker* channel EKO. We observed synaptic density reduction globally in the antennal lobe, compared with the control group where an inactive variant of tetanus toxin light-chain (TNT-) was introduced instead (Figure S5B, C). This confirmed the feasibility of using this transgenic reporter to measure synapses and the effect of neural activity on synaptogenesis.

We next used the pan-ORN *peb-GAL4* to knock down *beats/sides* that are expressed in ORNs and have known interacting partners. We also knocked down *side-VIII*, which is not expressed in ORNs, to serve as a presumable negative control (Figure 2A, B; Figure S2B). We found a pervasive synaptic reduction in *side* knockdowns compared to the control *RNAi,* but not *side-VIII* knockdowns (Figure 4J; Figure S5D, E). In contrast, knockdown of sparsely expressed genes, including *beat-IIa/b* and *side-IV*, didn’t lead to detectable differences from controls (Figure 4A; Figure S5D, E). Additionally, knockdown of other broadly expressed ones, including *beat-Ia*, *side-II, side-III*, and *side-VI*, also didn’t cause significant changes in synaptic density of ORNs, except *side-VII*, which increased the overall synaptic density in the antennal lobe (Figure 4A; Figure S5D, E). The discrepancy of loss-of-function effects between *side* and other broadly expressed ones could be due to the biochemical property of Side protein, which interacts with four Beat partners, more than any other Side family members (Figure 1A). It is likely that a lack of Side leads to a more substantial reduction in trans-synaptic adhesion than the lack of other Side proteins (Figure 4A). As opposed to *side* with potential functions in synapse induction, *side-VII* might play a role in inhibiting synapse formation. Though we were unable to map the *side-VII* expression pattern in the antennal lobe as the gene trap driver is unavailable, the single-cell RNA-seq datasets show that *side-VII* appears to be broadly expressed across ORN classes and may interact with its cognate partner Beat-IV, which is also broadly expressed across PN classes as well as ORN classes (Figures 2B, 3B; Figures S2B, S3B). Thus, Side-VII might inhibit synapse formation through either *trans* interaction with post-synaptic Beat-IV or *cis* binding with Beat-IV on the same membrane. Overall, our results suggest the bidirectional regulatory functions of Beat/Side proteins in ORN synapse formation.

Interestingly, analysis of previously reported antennal RNA-seq data (Deanhardt et al., 2023) from *Or47b* and *Or67d* mutants revealed differentially expressed *beat* and *side* genes (Figure S5F). We find that certain *beat* and *side* genes are up-regulated while others are down-regulated in olfactory receptor mutants (Figure S5F). These results suggest that these genes are responding to neural activity and might contribute to ORN-specific synapse development and maintenance. Notably, the transcriptional changes induced by OR-dependent neural activity, in addition to lineage factors, can contribute to the final combinatorial expression and synaptic function of Beat and Side proteins.

### Evolutionarily conserved expression of *beat/side* orthologs across insect ORNs

Next, we sought to interrogate the ORN class-specific expression of *beats/sides* in other species to gain some evolutionary insights into the regulation and function of these families of IgSF proteins. Recently reported transcriptome atlas of ORNs in two additional insect species, yellow fever mosquitoes *Aedes aegypti*, and clonal raider ants *Ooceraea biroi*, allowed us to perform comparative analyses of *beat/side* expression in the peripheral olfactory tissue at single-cell resolution. Yellow fever mosquito and clonal raider ants belong to the order Diptera and a distant order Hymenoptera, and diverged from the fruit fly *Drosophila melanogaster*, a Dipteran species, 260 million and 300 million years ago, respectively (Figure 5A). We first set out to identify the bona fide orthologs of *beats/sides* in mosquitoes and ants. To do this, we queried the protein sequence of *Drosophila* Beat-Ia and Side against the proteome database of two other species and ran multiple sequence alignment after selecting the hits with comparable lengths and similar AlphaFold-predicted structures (two Ig domains for Beats; five Ig domains and one Fibronectin domain for Sides, also see Materials and Methods, Figure S5A). We then built the phylogenetic tree of Beat/Side protein orthologs across the three species. This analysis yielded 13 *beat* genes and 10 *side* genes in the yellow fever mosquito, while the clonal raider ant has 10 *beats* and nine *sides* in its genome (Figure 5B, C). We observed several notable gene duplication/loss events in these species. For example, there is a single ant gene in the Beat-I clade, whereas there are three copies in mosquitoes and flies (Figure 5B). In contrast, three ant genes encode the Side-IV clade and three mosquito genes encode the Side-II clade, while only one fruit fly gene is in each clade (Figure 5C). Additionally, the *side-V* gene appears to be lost in clonal raider ants (Figure 5C). The phylogenetic tree indicates that: (1) gene duplication events generated ancestral *beat/side* paralogs earlier than insect species divergence; (2) additional gene duplication/loss events also occurred post-divergence of fruit fly, mosquito, and ant species.

**Figure 5.**
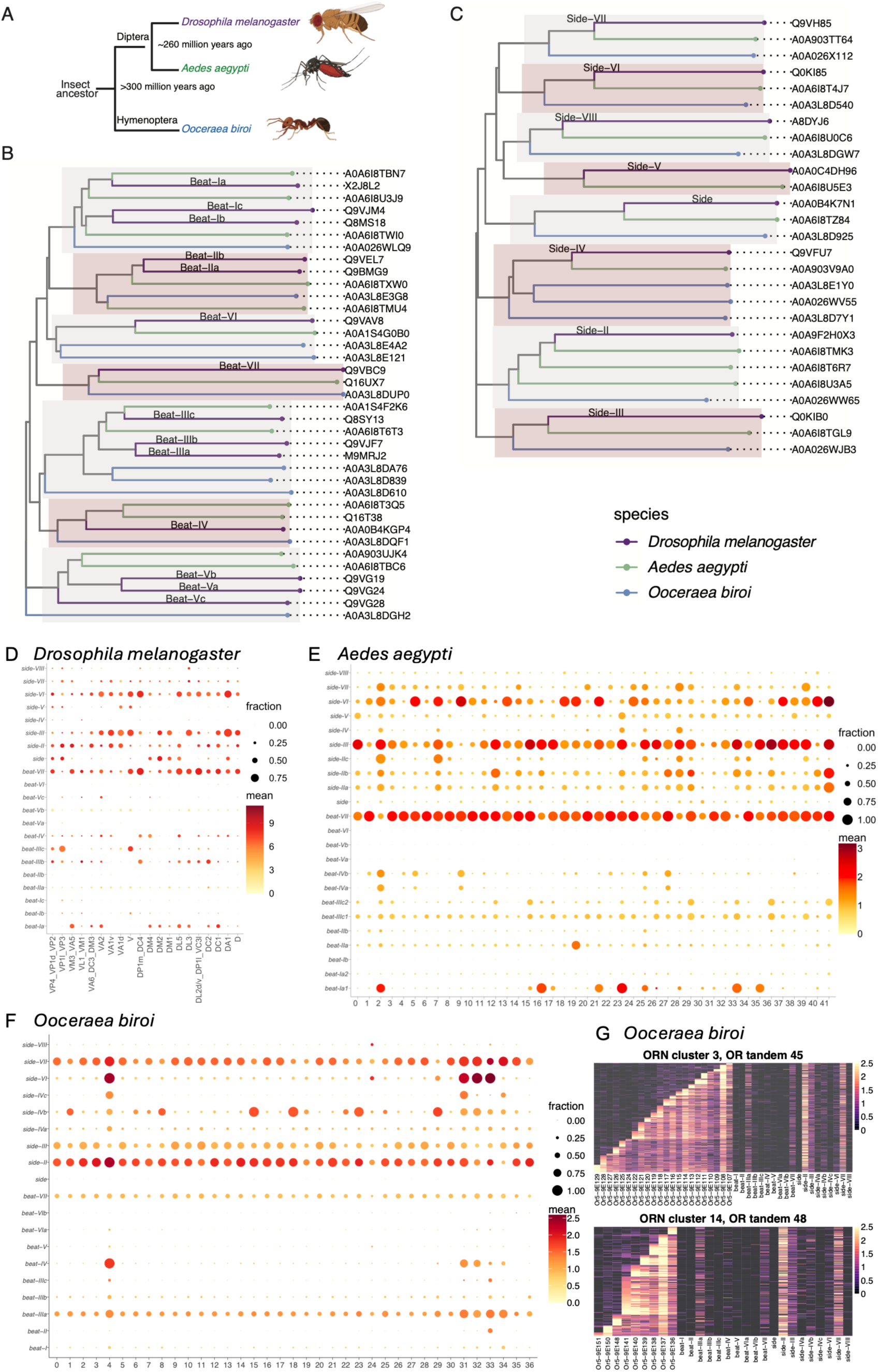
Evolutionarily conserved expression pattern of *beats/sides* across ORNs in insects. **(A)** Schematic phylogeny showing the evolutionary distance among three insect species. **(B)** Phylogeny tree of Beat orthologs in fruit flies *Drosophila melanogaster*, yellow fever mosquitoes *Aedes aegypti*, and clonal raider ants *Ooceraea biroi*. Each leaf is color-coded by the species, shown in the bottom right legend. Each ortholog is displayed as the UniProt protein ID, while the known *Drosophila* Beat names are labeled on the corresponding branch. **(C)** Phylogenetic tree of Side orthologs across three insect species. **(D)** Expression of *beats/sides* in the adult ORN classes of fruit flies based on the single-cell RNA-seq data from (McLaughlin et al., 2021). Each row is a gene, and each column represents an annotated ORN class. The size of each dot represents the percentage of positive cells in the given class (log2(CPM+1) > 0.5), and the color denotes the mean expression levels. This panel was reproduced from Figure S2B for comparison. **(E)** Expression of *beats/sides* in the adult ORN clusters of yellow mosquitoes based on the single-cell RNA-seq data from (Herre et al., 2022). The mosquito *beats/sides* were renamed based on their phylogenetic relations with the fruit fly orthologs. **(F)** Expression of *beats/sides* in the adult ORN clusters of clonal raider ants based on the single-cell RNA-seq data from (Brahma et al., 2023). The ant *beats/sides* were renamed based on their phylogenetic relations with the fruit fly orthologs. **(G)** Heatmap showing the expression of *beats/sides* and *OR* genes in two example ORN clusters of clonal raider ants, based on the single-cell RNA-seq data from (Brahma et al., 2023). *OR* genes are ordered from 5’ end to 3’ end in the tandem.

Based on the phylogenetic tree, we renamed the mosquito and ant genes according to their sequence similarities with the fruit fly orthologs (Table S1). We examined their expression in ORNs using published single-cell datasets (Brahma et al., 2023; Herre et al., 2022). Strikingly, we found combinatorial expression principles of *beat/side* genes in different ORN classes across the three insect species are conserved (Figure 5D, E, and F; Figure S6B): (1) genes that are not expressed in *Drosophila* ORNs, like *beat-Va/b/c*, *beat-VI*, and *side-VIII*, are also not expressed in ORNs of yellow fever mosquitoes (*beat-Va/b*, *beat-VI*, *side-VIII*) or clonal raider ants (*beat-V*, *beat-VIa/b*, *side-VIII*); (2) genes that are broadly expressed in *Drosophila* ORNs, like *beat-VII*, *side-II*, and *side-III*, are also broadly expressed in ORNs of yellow fever mosquitoes (*beat-VII*, *side-IIa/b*, and *side-III*) and clonal raider ants (*beat-VII*, *side-II*, and *side-III*); (3) genes that are expressed in a restricted pattern across *Drosophila* ORNs, like *beat-IIa/b*, beat*-IV*, and *side-IV* (Figure 2A, B), are also expressed in only a portion of ORN clusters in mosquitoes (*beat-IIa*, beat*-IVa/b*, and *side-IV*) and ants (*beat-II*, beat*-IV*, and *side-IVb/c*). Even though there is significant conservation of *beat/side* gene expression patterns across ORNs, there are also a few exceptions. For example, *beat-Ia*, expressed in many ORN classes in fruit flies and at high levels in restricted ORN classes in mosquitoes, is barely expressed in ant ORNs. In addition, *side* is expressed in many *Drosophila* ORN classes but is undetectable in mosquito and ant ORNs (Figure 5D, E, and F). These comparative analyses suggest that both the evolutionarily conserved and plastic *beat* and *side* genes in peripheral sensory neurons likely emerged due to genome evolution events that deleted or duplicated the members of the *beat* and *side* gene families and the regulatory sequences.

Interestingly, recent single-cell RNA-seq reports from clonal raider ant ORNs revealed regulatory differences in sensory receptor genes, where multiple *OR* transcripts are expressed in each ORN. In detail, ORNs that share a more similar transcriptome, and thus are clustered together in UMAP plots, usually express *ORs* located in a tandem array in the genome. In each ORN, a single start site is selected that transcribes *OR* genes downstream, while only the most upstream mRNA in the transcript is translated into a functional OR protein. This means the ORNs in each cluster are transcriptionally heterogeneous, containing multiple ORN classes defined by the functional expression of a singleton OR protein. To zoom in on ORN classes within each single-cell RNA-seq cluster, we plotted *beat/side* expression and the *ORs* of the tandem expressed in the cluster of each single cell as heatmaps (Figure 5G; Figure S6D). We found that each ORN class, defined by the expression of the most upstream *OR*, exhibits class-specific expression of *beat/sides*. For example, in cluster 3, the ORN class defined by the functional expression of Or5-9E109 doesn’t express the ant *side-IVb,* whereas other ORN classes functionally expressing the adjacent Ors, Or5-9E108, Or5-9E107, and Or5-9E110 appear to be positive for this gene (Figure 5G top). Similarly, ORN classes defined by the functional expression of Or5-9E151 or Or5-9E150 in cluster 14 don’t express the ant *beat-VII,* whereas many other ORN classes in this cluster do (Figure 5G bottom). This suggests, upon OR tandem choice, additional lineage mechanisms couple *OR* transcription initiation and *beat/side* expression specification in the clonal raider ant.

## Discussion

In this study, we comprehensively delineated the expression pattern of *beat/side* family genes in ORNs and PNs at single-cell and single-class levels. We found that each ORN or PN class exhibits a unique combinatorial expression of *beats/sides*, and this *beat/side* profile appears to be specified by the lineage-intrinsic genetic programs. In addition to analyzing the single-cell transcriptome of ORNs and PNs, we also generated *MiMIC*-based gene trap driver lines to probe the *beat/side* expression across ORNs and PNs *in vivo*. This approach enabled mapping the *beat/side* expression to additional ORN/PN types not covered in the single-cell RNA-seq datasets. Collectively, we found that *beat/side* profiles can diverge between some closely related ORNs or some closely related PNs, suggesting that the Beat/Side combination may add variability to cell surface codes, which biases their glomerular targeting. We also investigated one interacting pair, Beat-IIa and Side-IV, based on their matching expression pattern between partner ORNs and PNs, and found no apparent glomerular mistargeting in knockdowns of *Beat-IIa* in ORNs or *Side-IV* in PNs. Yet, knockdown of the *side* gene in ORNs resulted in diminished synapses in the antennal lobes, suggesting a role in synapse development. Moreover, we found evolutionarily conserved expression patterns and ORN-specific combinatorial signature principles for *beat* and *side* orthologs in the olfactory systems of mosquitoes and ants, suggesting the shared genetic programs establishing the *beat/side* profile. In sum, our studies implicate ORN/PN class-specific combinatorial Beat and Side protein expression and their roles in olfactory circuit assembly.

### Developmental regulation of *beat/side* expression in ORNs and PNs

We found the lineage-correlated expression profile of *beats/sides* in ORNs and PNs to be a general principle, suggesting the lineage-specific programs in setting the *beat/side* combinatorial expression. Our genetic labeling results and single-cell RNA-seq analyses also reveal the existence of both shared and divergent expression patterns of *beat/side* between closely related ORNs. We found that some sibling ORNs have different *beat/side* profiles and project to distant antennal lobe regions, while some maintain the putative “template” combinations found in many ORNs and project to neighboring or close glomeruli. Adjacently born PNs can also have shared or divergent *beat/side* expression profiles. We therefore propose that: (1) at the higher level of the developmental hierarchy, i.e., in ORN sensillar type decisions or PN lineage specification decisions, the *beat/*side “template” is set by lineage-specific factors; (2) the latest common precursor of ORNs (i.e. sensilla subtype precursors) or PNs may have diverse intrinsic factors that bias the expression of additional *beat/side* combinatorial expression while some do not; (3) such that at the final terminal selection decision, Beat/Side combinations may mediate the synapse programs in closely related ORNs or PNs. Furthermore, expression of additional cell surface proteins can increase the cell surface protein repertoire, further diversifying neuronal identification tags displayed on neurons utilized during circuit assembly. This two-step model, setting the lineage-specific cell surface molecular template, followed by within-lineage diversification, seems to be an efficient way to establish the cell type-specific surface molecular repertoire by genetically deterministic programs. This might be a developmental strategy to coordinate the combinatorial expression of duplicated paralogs.

### Molecular complexity and redundancy of Beats/Sides as cell surface “codes”

Side was first identified as the postsynaptic ligand, expressed in muscles, to attract motor neurons. Motor neurons expressing the receptor Beat-Ia follow the Side-labeled muscles and leave axonal fasciculations to innervate muscle targets (de Jong et al., 2005; Fambrough and Goodman, 1996; Pipes et al., 2001; Siebert et al., 2009; Sink et al., 2001). The embryonic expression pattern analysis expanding to other Beat/Side family paralogs also suggests that Beats are neuronal receptors for Sides expressed on peripheral tissues (Li et al., 2017b). For example, Side-VI is expressed in muscle fibers while its receptors, Beat-Vs, are expressed in motor neurons (Li et al., 2017b). This evidence jointly points to the fact that Sides are present primarily in postsynaptic sites, signaling to presynaptic Beats to promote synaptogenesis. Outside of the neuromuscular junction, recent reports in the fly visual circuits also suggest presynaptic Side-II and postsynaptic Beat-VI mediate synaptic recognition (Yoo et al., 2023). Additional studies revealed presynaptic (photoreceptor-expressed) Side-IV can induce synaptogenesis by acting as both ligands and receptors when interacting with postsynaptic Beat-IIa/b, forming synaptogenic complexes through their interactions with Beat-IIa/b and coreceptor Kirre (Osaka et al., 2024).

In the olfactory circuits consisting of ORNs and PNs, we found that *beat* and *side* genes are expressed in the same neurons. ORNs generally lack Beats but are abundant for Sides. This would point to a model where Sides primarily interact with PN-expressed Beats. However, ORNs do express a few Beats with their binding partner Sides as well as some orphan Beats. And PNs generally express both Sides and Beats. Similarly, DIPs and Dprs can also be expressed in the same cells, like adult ORNs (Barish et al., 2018) and larval motor/sensory neurons (Wang et al., 2022). There are several possible functional explanations based on Beat and Side interactions. First, Sides and Beats may have different subcellular localizations that position places of protein-protein interactions and cellular functions. For example, Beats might localize to PN dendrites to interact with Sides on ORN axon terminals, while Side proteins localized to PN axon terminals may interact with Beats expressed on the dendrites of third-order neurons. If this is the case, this binary subcellular preference may vary in different cell types to mediate interactions with other neurons. In motor neurons, Beats are preferentially sorted to axon terminals, whereas in the central nervous system or sensory neurons, Sides are preferentially sorted to axon terminals. Second, Beats/Sides can function in both dendrites and axons simultaneously and act as ligands and receptors bidirectionally. This points to a model in which the combinatorial code of multiple surface proteins establishes synaptic specificity recognition, likely by providing differential adhesive forces. Third, Beats and Sides in the same neuron might interact *in cis* on the same membrane in addition to their interactions *in trans* with Beat/Side proteins on other neurons. Interestingly, a recent study showed that DIP-Dpr interactions favor *cis* over *trans* when present in the same cell (Morano et al., 2025). It might also be the case that when Beats/Sides are localized together in the same membrane regions, it is the competition between *cis* and *trans* interactions with all present binding partners in the proximity that sets the “net” Beat/Side surface codes.

Additionally, combinations of cell surface adhesion molecules can form multi-molecular complexes, so the absence of any component may not abolish but just reduce the recognition efficacy or shift these complexes to new stoichiometric configurations (Honig and Shapiro, 2020; Sanes and Zipursky, 2020). Given this sophisticated context, it is not surprising that removing a single gene is insufficient to change glomerular organization. Regarding Beat/Side combinatorial complexity, deleting *side-IV* in PNs or *beat-IIa* in ORNs doesn’t ambiguate DM1 from adjacent glomeruli (Figure 4), and ORN-PN synaptic matching can still occur. Thus, the promiscuous and redundant functions of the Beat/Side family have the capacity to form an error-tolerant adhesion code, enabling recognition robustness. These are supported by our efforts to perturb a single *beat/side* gene that caused subtle to no defects (Figure S4A). Previously reported screens for CSMs mediating ORN-PN synaptic matching also didn’t report any defects in glomerular targeting with any *beat/side* knockdowns (Xie et al., 2019). A recent study has shown that collective manipulation of multiple cell surface molecules substantially rewires an ORN-PN circuit, while loss-of-function of a single one only causes minor mistargeting (Li et al., 2025; Lyu et al., 2025). Our expression dataset provides a roadmap to further crack the combinatorial coding nature of Beats and Sides by multiplexing perturbation.

Intriguingly, we observed weakened synapses in the antennal lobes when we knocked down the *side* gene, which suggests that Side and Beat protein interactions might mediate synapse formation in the olfactory circuits. This is supported by other recent studies in the visual system reporting Side-IV as a synapse-inducing protein (Osaka et al., 2024), and that Beat-VI expression gradients regulate synaptic density gradients through their interactions with Side-II (Dombrovski et al., 2025). These results suggest that Beat/Side interactions may inherently have synaptogenic functions and participate in the subglomerular synaptic organization during olfactory circuit development. Furthermore, since many *beats/sides* show persistent high expression levels from the pupal stage to adulthood, they might be recruited for synapse induction, formation, and stabilization, and modulated by OR signaling and/or neuronal activity.

### Evolutionarily conserved expression of *beats/sides* in ORNs of insects

We observed similar combinatorial expression patterns across ORN clusters among recently duplicated paralogs within the same species, as well as orthologs in ants and mosquitoes, suggesting evolutionarily conserved mechanisms that regulate their expression in ORNs. We speculate that ancestral paralogs of *beats/sides* with their regulatory elements acquired cell-type-specific expression patterns, which were generally retained during evolution and shared between species. More recent gene duplication events after speciation generated additional paralogs within each *beat/side* clade, where we observe both shared (*Drosophila beat-IIIa/b/c* and *beat-Va/b/c*, *A. aegypti beat-IVa/b* and *side-IIa/b*, *O. biroi beat-VIa/b*) and divergent expression patterns (*A. aegypti beat-Ia1/a2* and *side-IIb/c*, O*. biroi side-IVa/b/c*). Though some *beat* and *side* genes exhibit evolutionary plasticity in their expression patterns, conservation of the expression of many *beat/side* genes in ORNs appears to be a general rule. This implies that there are selection pressures on the operative roles that these genes play in the ORN development and function.

Interestingly, in clonal raider ants, there appear to be two steps specifying the ORN fate: first, choosing the OR tandem and second, choosing the OR transcription start site within the tandem. Our results suggest that the *beat/side* ortholog expression is also coupled with these two steps. *Drosophila* appears to exploit the genetically deterministic strategy to specify the combinatorial expression of cell surface proteins to organize diverse ORN classes into the hardwired circuits. However, when neuronal types become more heterogeneous, possibly exceeding a diversity limit, such a solely genetics-dependent program may be incapable. This is likely to be the case in the olfactory systems of ants with many more classes of ORNs.

Recent studies suggest ORs themselves may play instructional roles in glomerular targeting in ants, different from fruit flies but similar to mammals (Duan and Volkan, 2020; Ryba et al., 2020; Trible et al., 2017; Yan et al., 2017). It is possible that the ORN class-specific expression of *beat/side* is also regulated by ORs in ants.

### Limitations of the study

Many *beat/side* genes transcribe multiple splice isoforms, which adds an additional layer of molecular diversity. This study didn’t examine the expression of *beat* and *side* splice isoforms in the olfactory system. Our transgenic *GAL4* driver lines generally trap the common regions across isoforms and cannot address isoform-specific expression. Single-cell RNA-seq approaches that resolve *beat* and *side* splice isoforms or expression analysis of isoform-specific *GAL4*s would be needed to this end.

This study only reveals the *beat* and *side* expression at the transcriptional level and does not address the Beat and Side protein-level distributions. Some studies reported large discrepancies between transcripts encoding surface molecules and their protein abundance in developing PNs (Li et al., 2020b), suggesting regulation at the level of translation, protein stability, modification, sorting, and delivery to the cell surface.

### Future directions

In the future, it will be interesting to investigate how regulatory sequence evolution contributes to the conserved and divergent expression patterns of *beat* and *side* paralogs across species. Additional biochemical characterization of Beats/Sides protein-protein interactions in different insect species will provide novel insights into the co-evolution of coding sequences with regulatory sequences to reconcile their functional evolution in circuit assembly.

Understanding the biochemical features of Beat and Side proteins will greatly inform the functional analysis in the circuit assembly context. This includes developing a more sensitive and specific test to deorphanize Beat-Side interactions, characterizing whether Beats/Sides are transmembrane proteins or membrane-anchored like DIPs/Dprs (Lobb-Rabe et al., 2024), and testing whether they interact *in trans* and/or *in cis*. If they are transmembrane proteins, then how are their functions mediated by intracellular and extracellular protein domains? Proximity labeling-based proteome profiling, like BIO-ID, can identify these protein interactors, which will inform further functional investigation. A recent study reported that the N-terminal domain of Beat-Ia is localized to the cell surface while the C-terminus accumulates in the nucleus; this implies that Beat-Ia might undergo proteolytic cleavage for proper function (Heymann et al., 2022). Future biochemical, genetic, and neurophysiological studies will help reveal the function of Beat and Side proteins in recognition specificity and synaptic transmission.

---

**Figure S1.**
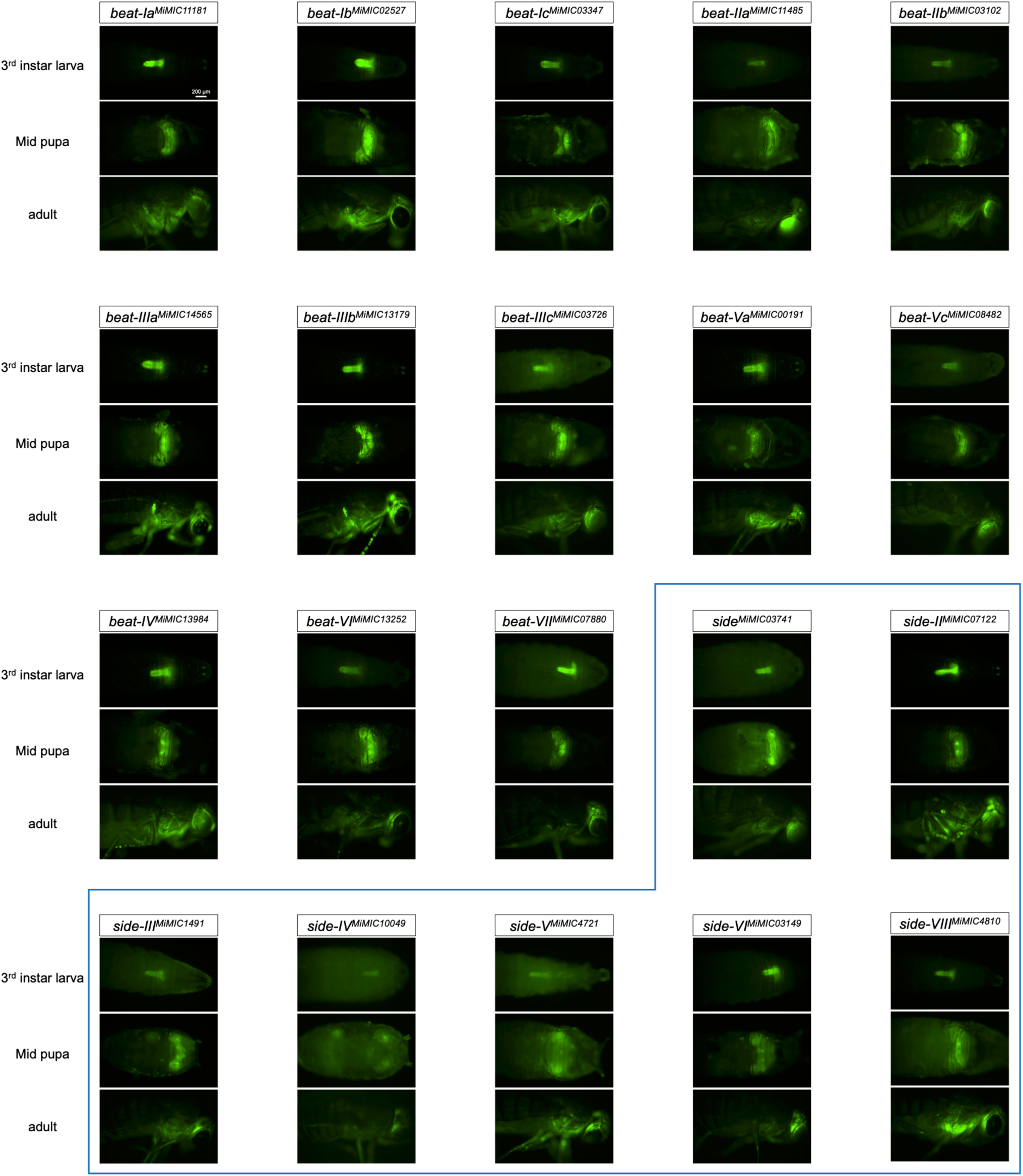
*beat* and *side* expression are enriched in the nervous system throughout development revealed by *MiMIC*-based gene trap drivers. Related to Figure 1. Whole-animal fluorescence images of each *beat/side* gene trap GAL4 driving myr.GFP in the 3^rd^ instar larval stage, mid-pupal stage (∼48h APF), and adult.

**Figure S2.**
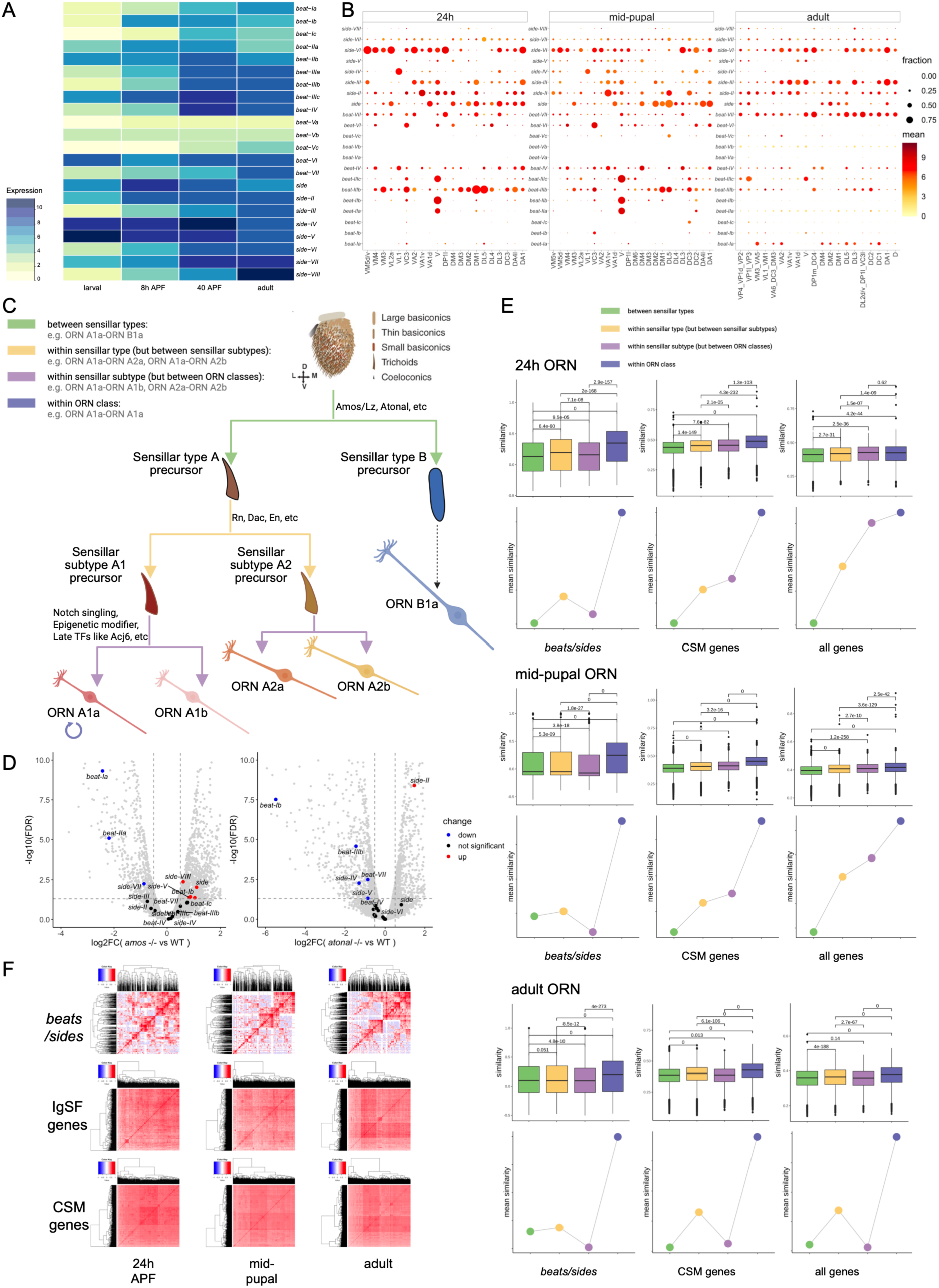
Characterization of *beat/side* expression in the published datasets of bulk antenna and single-cell RNA-seq of ORNs. Related to Figure 2. **(A)** Dynamic *beats/side* expression in antennal tissues over development. Heatmap showing the temporal dynamic expression of *beats/sides* from 3^rd^ instar larval antennal discs (larval), 8h APF pupal antennal discs, 40h APF antennae, and adult antennae. The color indicates the log-scaled relative expression values (base mean) of each gene at each stage from DESeq results. **(B)** *beat/side* genes are differentially expressed across ORN classes at single-cell resolution. Each column is an annotated ORN class, and each dot represents the expression level of the row gene in the given ORN class, summarizing from the single-cell (24h APF, mid-pupal stage) or single-nucleus RNA-seq datasets (adult). The size of the dot denotes the fraction of “positive” cells in the given ORN class (log2(CPM+1) > 0.5), and the color denotes the mean expression of the positive cells. **(C)** Hierarchical genetic programs control ORN lineage and glomerular targeting. A simplified model illustrating the ORN-ORN kinship. ORNs are housed in sensillar subtypes within each sensillar type, representing different genetic lineages. ORNs mapped to the same class are defined as “within ORN class”, like ORN A1a and ORN A1a; ORNs of different classes but housed in the same sensillar subtype are defined as “within sensillar subtype”, like ORN A1a and ORN A1b; ORNs from different sensillar subtypes but belonging to the same sensillar type, are defined as “within sensillar type”, like ORN A1a and ORN A2a; and the furthest kinship is ORNs housed in distinct sensillar types, i.e., “between sensillar types”, like ORN A1a and ORN B1a. Schematic was generated by BioRender. The antenna structure is reproduced from (Li et al., 2013). **(D)** *beat/side* expression in the antenna is regulated by sensillar type-specification factors *amos* and *atonal.* Volcano plot showing the differentially expressed genes in *amos* mutant antennae compared with wild-type antennae (left panel) or *atonal* mutant antennae compared with wild-type antennae (right panel). Significantly downregulated *beat/side* genes are colored in blue, and significantly upregulated *beat/side* genes are colored in red. Significance is determined by FDR < 0.05 from EdgeR results. The horizontal dashed line in each plot is FDR = 0.05. Two vertical dashed lines in each plot are log2FC = -0.5 and 0.5, respectively. Gray dots are all other genes detected. **(E)** Pairwise correlation between ORNs at three stages reveals the *beat/side* combinatorial expression is correlated with the ORN kinship. Top row in each stage: boxplot showing the similarity measured by Spearman’s correlation between two cells from the indicated stage. The pairwise relation is categorized into four groups depending on the annotated ORN class and the corresponding sensillar type/subtype lineage. Bottom row in each stage: each dot is the mean similarity of the corresponding category in the top boxplot. The Spearman’s correlation was calculated based on the expression of all beat/side genes, all CSM-encoding genes, or the whole transcriptome (all genes). P values are from Mann–Whitney U tests without multiple comparison adjustments. Figure 2E is reproduced here for comparison. **(F)** *beat/side* combinatorial expression is more cell-population specific than pan-IgSF or pan-CSM genes across ORNs. Heatmap showing the Spearman’s correlation of combinatorial gene expression between the row cell and the column cell across three developmental stages (24h APF, mid-pupal, and adult). The correlation was computed based on the combinatorial expression of *beat-side* genes, IgSF-encoding genes, or CSM-encoding genes. Cells are hierarchically clustered, as shown in the phylogeny tree in each heatmap. Correlation is shown as a spectrum from blue (-1) to red (1). The diagonal is the similarity of each cell to itself, and thus the correlation always equals 1.

**Figure S3.**
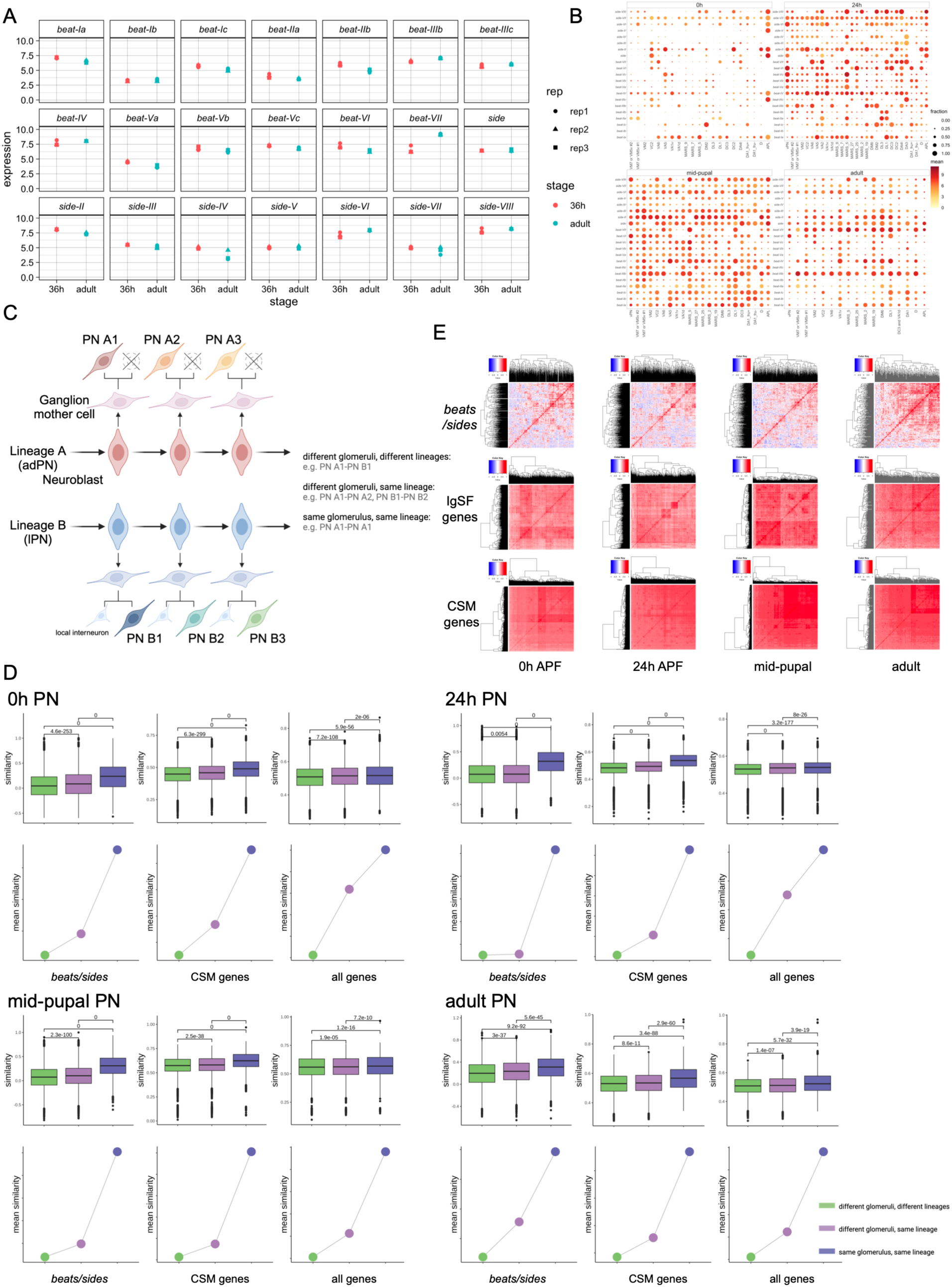
Characterization of *beat/side* expression in the published datasets of bulk and single-cell RNA-seq of PNs. Related to Figure 3. **(A)** *beat/side* expression in bulk PNs over development. Point plots showing the *beat/side* expression in the sorted bulk PNs from 36h APF and the adult stage. The shape of each point denotes the biological replicate. The expression value is log2(CPM+1). **(B)** *beat/side* genes are differentially expressed across PN classes at single-cell resolution. Each column is an annotated PN class, and each dot represents the expression level of the row gene in the given PN class, summarizing from the single-cell RNA-seq datasets. The size of the dot denotes the fraction of “positive” cells in the given PN class (log2(CPM+1) > 0.5), and the color denotes the mean expression of the positive cells. APL, anterior paired lateral neurons. APL and vPN clusters were not used for the correlation analysis in D. **(C)** Sequential genetic programs control PN lineage and glomerular targeting. We only used PNs mapped to adPN and lPN lineage for correlation analysis. PNs from these two lineages are excitatory and project to a single glomerulus, whereas the third lineage, vPN neurons, which were removed for downstream analysis, are inhibitory GABAergic and can project to multiple glomeruli (Liang et al., 2013; Marin et al., 2002). In the adPN lineage, one of the two post-mitotic neurons survives and develops into a PN. In the lPN lineage, both post-mitotic neurons survive, one becoming a PN and the other becoming a local interneuron (Lin et al., 2012; Lin et al., 2010). Schematic was generated by BioRender. **(D)** Pairwise correlation between PNs at four stages reveals the *beat/side* combinatorial expression is correlated with the PN kinship. Top row in each stage: boxplot showing the similarity measured by Spearman’s correlation between two cells from the indicated stage. The pairwise relation is categorized into three groups depending on the annotated PN class identity and the corresponding lineage identity. Bottom row in each stage: each dot is the mean similarity of the corresponding category in the top boxplot. The Spearman’s correlation was calculated based on the expression of all beat/side genes, all cell surface molecule (CSM)-encoding genes, or the whole transcriptome (all genes). P values are from Mann–Whitney U tests without multiple comparison adjustments. Figure 3E is reproduced here for comparison. **(E)** *beat/side* combinatorial expression is more cell-population specific than pan-IgSF or pan-CSM genes across PNs. Heatmap showing the Spearman’s correlation of combinatorial gene expression between the row cell and the column cell across four developmental stages (0h, 24h APF, mid-pupal, and adult). The correlation was computed based on the combinatorial expression of *beat-side* genes, IgSF-encoding genes, or cell surface molecule-encoding genes. Correlation is shown as a spectrum from blue (-1) to red (1). Cells are hierarchically clustered, as shown in the phylogeny tree in each heatmap. The diagonal is the similarity of each cell to itself, and thus the correlation always equals 1.

**Figure S4.**
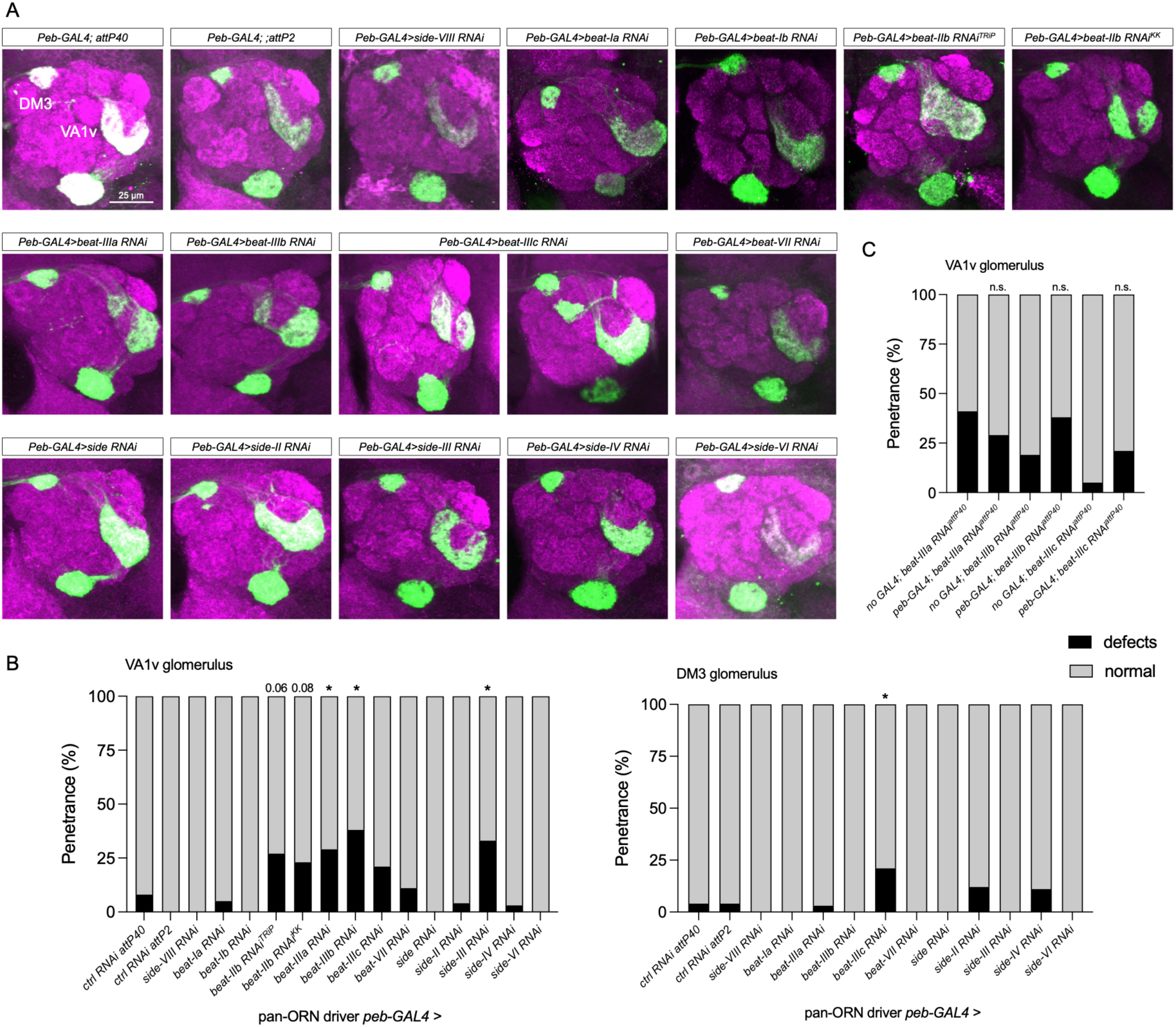
Perturbation of *beats/sides* in ORNs led to no or minor defects in glomerular organization. Related to Figure 4. **(A)** Representative antennal lobes of controls and knockdowns of *beat/side* genes. Knockdown of *beat-IV* or *side-V* with this *peb-GAL4* driver caused lethality, and thus the data were missing. **(B)** Quantification of the percentage of lobes showing glomerular defects of VA1v and DM3 glomeruli in each condition. N=20-50 antennal lobes examined. P values were determined by comparison to control groups. *, p<0.05. **(C)** Quantification of the percentage of lobes showing glomerular defects of VA1v glomerulus of *beat-IIIa/b/c* perturbation with additional *RNAi* transgenic background controls. N=22-34 antennal lobes. n.s., not significant.

**Figure S5.**
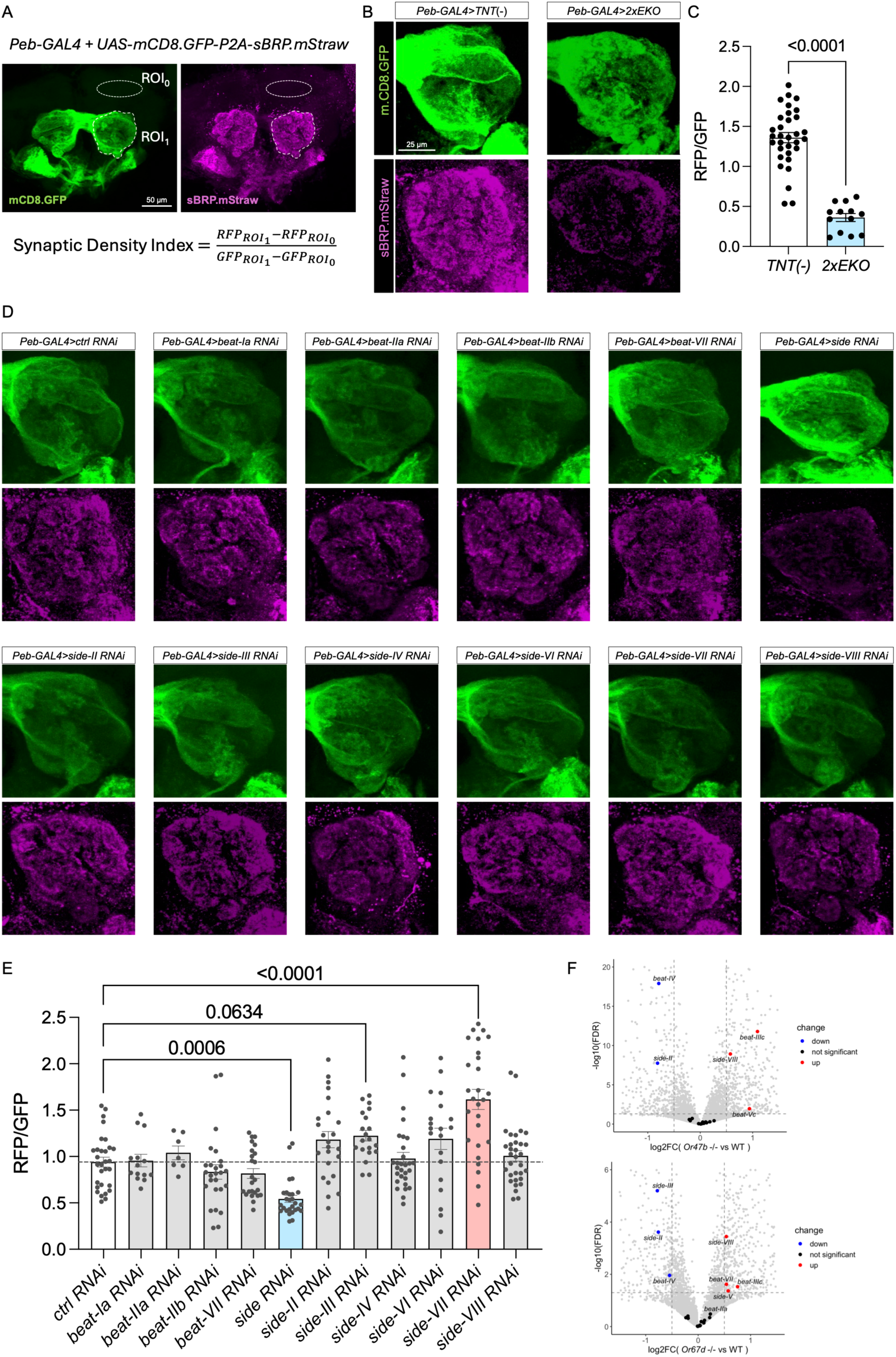
The effects of *beat/side* perturbation on synaptic density in ORNs. Related to Figure 4. **(A)** Synaptic density is quantified by the ratio of the average fluorescence intensity of RFP and GFP in the region of interest, i.e., the antennal lobe. Synaptic active zone marker, short BRP protein.mStraw, and membrane marker, mCD8.GFP, are translated from the single transcript but localized in different subcellular regions (axon terminals vs general neurites). **(B)** Representative antennal lobes showing the effects of blocking neuronal activity of ORNs on gross glomerular synaptic density, visualized by the membrane marker GFP and presynaptic marker RFP. Scale bar representing 25 µm also applies to each panel in (D). **(C)** Quantification of (B). Each dot is a single antennal lobe. N = 43 and 13 antennal lobes, respectively. P-value from the unpaired t-test between groups is shown. **(D)** Representative antennal lobes showing the effects of knockdown of *beats/sides* on gross glomerular synaptic density, visualized by the membrane marker GFP and presynaptic marker RFP. Knockdown of *beat-IV* or *side-V* with this *peb-GAL4* driver caused lethality, and thus the data were missing. There is little to no expression of *beat-Ib/c*, *beat-Va/b/c*, or *beat-VI* across ORNs; *beat-IIIa/b/c*, though expressed broadly, are orphans. These genes were not tested for their loss-of-function effects on synaptic density. **(E)** Quantification of (D). Each dot is a single antennal lobe. N = 30, 14, 7, 26, 24, 27, 24, 20, 29, 20, 27, 32 antennal lobes, from left to right. Only P-values that are significant or near the edge from multiple comparisons post-ordinary-one-way ANOVA are shown. The dashed horizontal line indicates the mean values in the *ctrl RNAi* group. Groups of *ctrl RNAi* and *side RNAi* are the same samples as in Figure 4J. **(F)** Volcano plot showing the differentially expressed genes in *Or47b* mutant antennae (top) and *Or67d* mutant antennae (bottom) compared with wild-type antennae. Significantly downregulated *beat/side* genes are colored in blue, and significantly upregulated *beat/side* genes are colored in red. Significance is determined by FDR<0.05 from DESeq2 results. The horizontal dashed line in each plot is FDR = 0.05. Two vertical dashed lines in each plot are log2FC = -0.5 and 0.5, respectively. Gray dots are all other genes detected. Data were reanalyzed from (Deanhardt et al., 2023).

**Figure S6.**
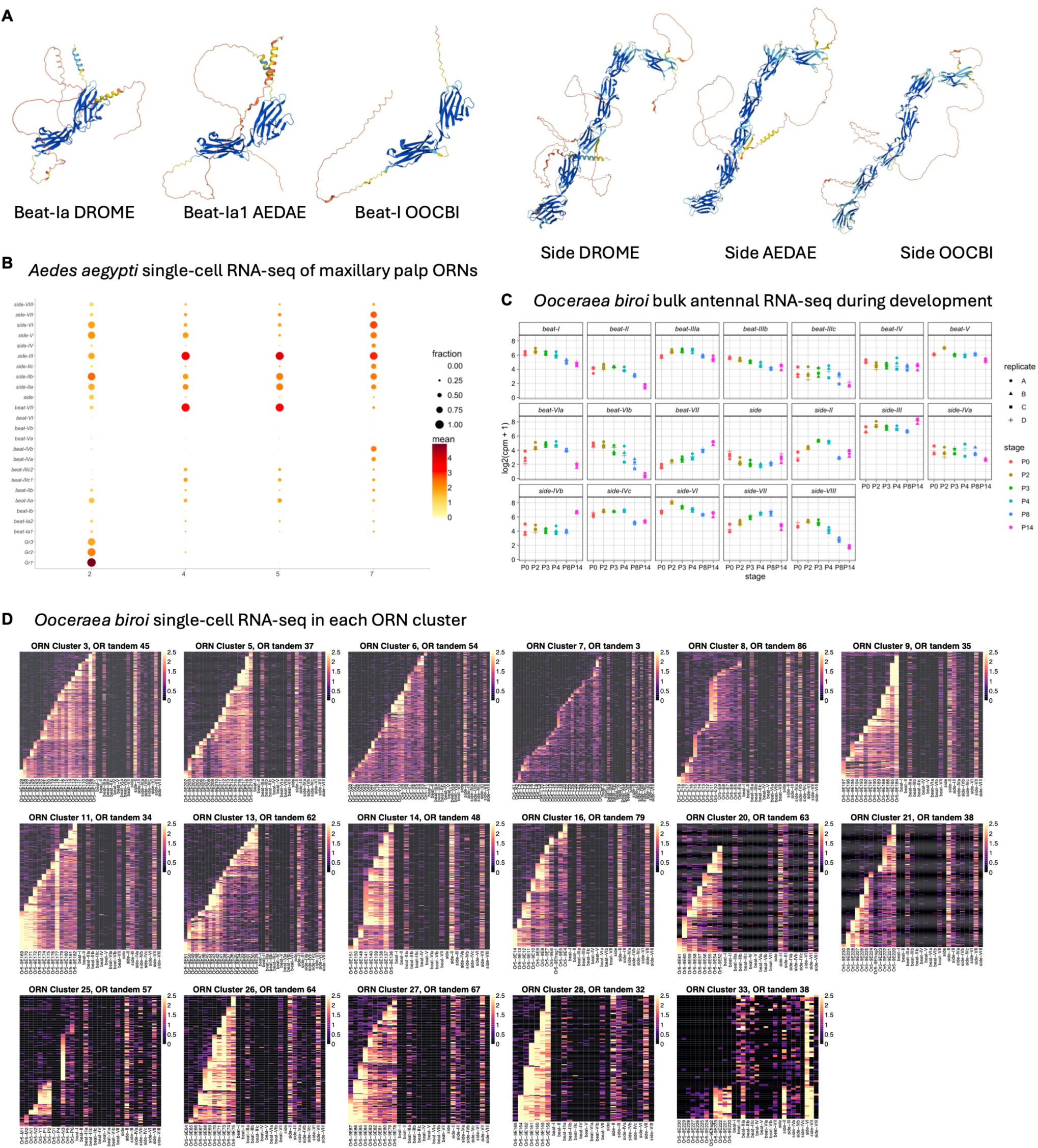
Additional analyses of *beat/sides* expression in mosquitoes and ants. Related to Figure 5. **(A)** AlphaFold-predicted structures of example Beat and Side orthologs in fruit flies *Drosophila* melanogaster (DROME), yellow fever mosquitoes *Aedes aegypti* (AEDAE), and clonal raider ants *Ooceraea biroi* (OOCBI). **(B)** Expression of *beats/sides* in the adult maxillary palp-housed ORN clusters of yellow mosquitoes based on the previously published single-cell RNA-seq data (Herre et al., 2022). **(C)** Bulk RNA-seq showing the expression of *beats/sides* in developing antennal tissues of clonal raider ants, reanalyzed from the data previously published (Ryba et al., 2020). **(D)** Heatmap showing the expression of *beats/sides* and *OR* genes in all ORN clusters of clonal raider ants that exhibit the ladder expression pattern of *OR* genes from 5’ end to 3’ end along a tandem, based on the single-cell RNA-seq data from (Brahma et al., 2023). *OR* genes are ordered from 5’ end to 3’ end in tandem in each cluster. Two clusters shown in Figure 5G are also reproduced here for comparison. Data reanalyzed from (Brahma et al., 2023).

## Materials and Methods

### *Drosophila* stocks and genetics

Flies were raised in classic molasses media provided by Archon Scientific. Most crosses were kept at room temperature (23 °C), except the RNAi experiments performed at 28 °C to maximize the knockdown efficiency. For RNAi experiments, male and virgin female flies were mixed at room temperature for three days to facilitate mating. Then flies were raised at a 28 °C incubator until 5-7 days after eclosion before dissection. The transgenic *Drosophila melanogaster* strains used in this study are listed in Table 1. BDSC, Bloomington Drosophila Stock Center; VDRC, Vienna Drosophila Resource Center.

**Table 1.** References and identifiers of the transgenic flies used in this study.

| **Genotype** | **Reference** | **Identifier** |
| --- | --- | --- |
| *hs-cre, vas-phiC31* (chrX) | (Diao et al., 2015) | BDSC 60299 |
| *Trojan-Gal4.0; Dr/TM3* | (Diao et al., 2015) | BDSC 60301 |
| *Sp/CyO; Trojan-Gal4.0* | (Diao et al., 2015) | BDSC 60302 |
| *Trojan-Gal4.1; Dr/TM3* | (Diao et al., 2015) | BDSC 60304 |
| *Sp/CyO; Trojan-Gal4.1* | (Diao et al., 2015) | BDSC 60305 |
| *Trojan-Gal4.2; Dr/TM3* | (Diao et al., 2015) | BDSC 60307 |
| *Sp/CyO; Trojan-Gal4.2* | (Diao et al., 2015) | BDSC 60308 |
| *10XUAS-IVS-myr.GFP attP40* |  | BDSC 32198 |
| *10XUAS-IVS-myr.GFP attP2* |  | BDSC 32197 |
| *beat-Ia-GAL4 [MI11181] chr2* | This study | N.A. |
| *beat-Ib-T2A-GAL4 [MI02527] chr2* | This study | N.A. |
| *beat-Ic-GAL4 [MI03347] chr2* | This study | N.A. |
| *beat-Ic-T2A-GAL4 [MI01467] chr2* | This study | N.A. |
| *beat-IIa-T2A-GAL4 [MI11485] chr3* | (Lee et al., 2018) | BDSC 76215 |
| *beat-IIb-T2A-GAL4 [MI03102] chr3* | This study | N.A. |
| *beat-IIIa-T2A-GAL4 [MI14565] chr2* | This study | N.A. |
| *beat-IIIb-T2A-GAL4 [MI13179] chr2* | (Lee et al., 2018) | BDSC 76227 |
| *beat-IIIc-T2A-GAL4 [MI03726] chr2* | This study | N.A. |
| *beat-IV-T2A-GAL4 [MI13984] chr3* | This study | N.A. |
| *beat-Va-T2A-GAL4 [MI00191] chr3* | This study | N.A. |
| *beat-Vc-T2A-GAL4 [MI08482] chr3* | This study | N.A. |
| *beat-VI-GAL4 [MI13252] chr3* | This study | N.A. |
| *beat-VII-GAL4 [MI07880] chr3* | This study | N.A. |
| *side-T2A-GAL4 [MI03741] chr3* | This study | N.A. |
| *side-II-T2A-GAL4 [MI07122] chr2* | This study | N.A. |
| *side-III-GAL4 [MI01491] chr3* | This study | N.A. |
| *side-IV-GAL4 [MI10049] chr3* | This study | N.A. |
| *side-V-GAL4 [MI04721] chr2* | This study | N.A. |
| *side-VI-T2A-GAL4 [MI03149] chr3* | This study | N.A. |
| *side-VII-GAL4 [MB10368] chr3* |  | BDSC 29116 |
| *side-VIII-T2A-GAL4 [MI04810] chr2* | This study | N.A. |
| *ey-FLP* (chrX) | (Newsome et al., 2000) | BDSC 5580 |
| *GH146-FLP* (chr2) | (Hong et al., 2009) | BDSC 81291 |
| *UAS>STOP>mCD8.GFP* (chr2) | (Hong et al., 2009) | BDSC 30125 |
| *UAS>STOP>mCD8.GFP* (chr3) | (Hong et al., 2009) | BDSC 30032 |
| *peb-GAL4* (chrX) | (Sweeney et al., 2007) | N.A. |
| *GH146-GAL4* (chrX) |  | BDSC 91812 |
| *UAS-Dcr* (chrX) |  | BDSC 58726 |
| *Or42b-mCD8.GFP* (chr2) |  | BDSC 52648 |
| *Or92a-mCD8.GFP* (chr2) |  | BDSC 52646 |
| *Or47a-syt.GFP, Or47b-syt.GFP, Gr21a-syt.GFP* (chr2) | Volkan lab stock | N.A. |
| *UAS-Luc RNAi attP2* |  | BDSC 31603 |
| *attP40* |  | BDSC 36304 |
| *UAS-beat-Ia RNAi attP40* |  | BDSC 64938 |
| *UAS-beat-Ib RNAi attP40* |  | BDSC 57157 |
| *UAS-beat-IIa RNAi attP2* |  | BDSC 28072 |
| *UAS-beat-IIb RNAi attP40* |  | BDSC 57157 |
| *UAS-beat-IIb RNAi KK* (chr2) |  | VDRC 104935 |
| *UAS-beat-IIIa RNAi attP40* |  | BDSC 64526 |
| *UAS-beat-IIIb RNAi attP40* |  | BDSC 56984 |
| *UAS-beat-IIIc RNAi attP40* |  | BDSC 50941 |
| *UAS-beat-IV RNAi attP40* |  | BDSC 56981 |
| *UAS-beat-VII RNAi attP40* |  | BDSC 60056 |
| *UAS-side RNAi attP2* |  | BDSC 50642 |
| *UAS-side-II RNAi KK* (chr2) |  | VDRC 107512 |
| *UAS-side-III RNAi KK* (chr2) |  | VDRC 103669 |
| *UAS-side-IV RNAi GD* (chr2) |  | VDRC 16636 |
| *UAS-side-V RNAi attP40* |  | BDSC 61953 |
| *UAS-side-VI RNAi KK* (chr2) |  | VDRC 103456 |
| *UAS-side-VIII RNAi attP40* |  | BDSC 62897 |
| *UAS-SynLight attP2* | (Aimino et al., 2023) | BDSC 602367 |
| *UAS-TNT (-)* (chr2) | (Sweeney et al., 1995) | BDSC 28840 |
| *2xUAS-EKO* (chr2) | (White et al., 2001) | BDSC 40974 |

### Generating *T2A-GAL4* transgenic flies from *MiMIC* lines

We used the *in vivo* genetic cross-based method (Diao et al., 2015) to swap the *T2A-GAL4* construct with the *MiMIC* cassette. Briefly, by multiple crosses, the dual-recombinase (*hs-cre* + *vas-phiC31*) helper component, the in-frame *T2A-GAL4* donor component, and the target *MiMIC* locus in the genes of interest were introduced together into one parental animal, where the *T2A-GAL4* donor sequence was excised out by Cre recombinase and inserted into the *MiMIC* docking site by germline-expressed phiC31 integrase. When this parental animal was crossed to the *UAS-myr.GFP* reporter line, the recombinant progeny would have GFP expression driven by the *MiMIC-GAL4* if *T2A-GAL4* was inserted in the correct frame and orientation. The recombinant was crossed with a double-balancer line to make a stable stock. Specifically, we also found that the *T2A-GAL4* construct could be used in any 5’ UTR-located *MiMIC* sites in a frame-independent manner. In this case, the *GAL4* coding sequence with ATG start codon in the 5’ UTR would hijack the translation of the native gene as an upstream open reading frame, and the *linker-T2A* sequence could be neglected. As the *GAL4* open reading frame is within the whole transcript of the native gene, it is still expected to faithfully represent the transcription levels and patterns of the native gene. We name the driver line *gene-GAL4* if *GAL4* is within 5’ UTR of the host gene (Table 1).

### Whole animal fluorescence imaging of *beat/side* expression

The whole animal fluorescence imaging was performed on an Olympus BX51WI upright scope equipped with a C11440-36U camera. Larvae and adults were killed by 70% ethanol first and mounted on slides without coverslips. Pupae were directly mounted on slides without coverslips. Larvae and pupae were dorsal side up, whereas adults were mounted with the lateral side up. Images were acquired under blue light and the GFP channel, with manually adjusted exposure time.

### Immunohistochemistry

Flies were first killed with 70% ethanol. Then, brains were dissected in PBST buffer (0.2% Triton X-100 in 1X PBS), fixed in 200 μL centrifuge tubes with 4% paraformaldehyde for 30 minutes, followed by three 10-minute washes in PBST. Primary antibody mix (150 μL) was then added to the tubes, incubating brains on the orbital shaker at 4 °C overnight. Brains were washed three times for 20 minutes each with PBST at room temperature before secondary antibody staining. Secondary antibody mix (150 μL) was then added to the tubes, incubating brains on the orbital shaker at 4 °C overnight. Brains were washed three times for 20 minutes each with PBST at room temperature again before being mounted on the imaging slides. Natural goat serum (1%) was added to the primary and secondary antibody mix for blocking. The following primary antibodies with dilution ratio were used: rabbit anti-GFP (Invitrogen, 1:1000), rat anti-Ncad (DSHB, 1:20), rabbit anti-DsRed (TaKaRa Bio, 1:250); the following secondary antibodies with dilution ratio were used: Alexa Fluor 488 goat anti-rabbit IgG (Invitrogen, 1:1000), Alexa Fluor 647 goat anti-rat IgG (Invitrogen, 1:200), Alexa Fluor Cy3 goat anti-rabbit IgG (Invitrogen, 1:200). Both primary and secondary antibody cocktails were diluted in PBST.

Specifically, for Brp synaptic analysis, we only stained RFP mStraw and Ncad without staining GFP for two reasons. First, the transgenic mStraw fluorescence of the synaptic marker is very low and needs a very high laser power to detect the signal, while the natural GFP fluorescence is bright enough. Second, as we later normalized the RFP intensity by the GFP intensity as the proxy for synaptic density in the region of interest, no staining of GFP can reduce the variability introduced during immunohistochemistry.

### Confocal Imaging

To prepare brain samples for imaging, a mounting solution (Flouromount-G, SouthernBiotech) was added to the brains on the glass slide before the cover slip was mounted. Confocal images were acquired by either an Olympus Fluoview FV1000 microscope or a Zeiss 880 microscope with a 40X or 60X objective lens. Brains were scanned through the Z-axis from the posterior side to the most anterior side of the antennal lobes. For *beat/side-GAL4*-based glomerular innervation analysis, imaging parameters varied between different *GAL4* drivers to highlight the lowly expressing glomeruli. For phenotypical determination, imaging parameters were kept consistent between experimental and control groups.

### *beat/side* glomerular expression pattern analysis of *beats/sides*

For *ey-FLP* induced labeling of each *beat/side*-expressing ORN class, we found *ey-FLP* expression appears leaky in germline and crossing *ey-FLP* first with *UAS>STOP>mCD8.GFP* may lead to the removal of the *STOP* sequence constitutively and thereafter fail to restrict the GAL4-driven GFP expression solely in ORNs. To overcome this issue, we first crossed each *GAL4* driver line to the *ey-FLP* line and then crossed to *UAS>STOP>mCD8.GFP* reporter line.

For *GH146-FLP* induced labeling of each *beat/side*-expressing PN class, each *GAL4* driver line can be directly crossed to the *GH146-FLP*; *UAS>STOP>mCD8.GFP* line. Only the results from female animals were reported in this study.

As we aimed to determine the ORN or PN expression pattern of *beats/sides* in a binary manner, i.e., positive or negative, we didn’t use the constant parameters to acquire the images for different *GAL4* driver lines. We adjusted the acquisition parameters in order to clearly show even lowly expressing glomeruli. After scanning each brain from the most posterior end to the most anterior end of the antennal lobe, we referred to the previously characterized glomerular map (Task et al., 2022) and manually determined the glomerular identity and the corresponding *beat/side* expression, assigning 0 to negative expression and 1 to positive expression. At least three brains were examined for each *GAL4* driver line. Notably, for the *GH146-FLP-*based labeling, we occasionally found inconsistent glomerular innervation patterns between individuals and even right and left lobes of the same brain (data not shown). This is likely due to the incomplete excision of the *STOP* cassette in a subset of PNs. As each glomerulus could be innervated by as few as one PN, it is possible that some glomeruli are not labeled in an antennal lobe. We thus determined these glomeruli to be positive if we observed the GFP signal for that glomerulus in two or more antennal lobes. The expression matrix, expression value (0,1) of each gene in each glomerulus, was then input to the downstream clustering analysis.

### Phenotype quantification

The measurement of the glomerular phenotype was determined by the proportion of antennal lobes showing morphological or positional abnormalities across all studied brains within each group, compared to the controls. The P-value was determined using the two-tailed Fisher’s exact test, using the integrated features of the GraphPad Prism 9 software.

To quantify synaptic density at the gross antennal lobe level, we first obtained the Z-stack of confocal sections by maximum projection. Then, we manually selected the regions of interest (ROI) according to the Ncad channel, which are the right and left antennal lobes. We also selected a control region in the dorsal part of the brain where the driver *peb-GAL4* has no expression. We then subtracted the average intensity of the RFP channel of the control region from that of the ROI. This is the adjusted mean RFP intensity of the ROI. We did the same thing to get the adjusted mean GFP intensity of the ROI. We finally calculated the ratio between the adjusted mean intensity of RFP and the adjusted mean intensity of GFP as the readout for each antennal lobe. An unpaired t-test or multiple comparisons post ordinary one-way ANOVA was used to compare the average RFP/GFP ratio between groups.

### Bulk tissue RNA-seq and single-cell RNA-seq analysis from previously published datasets for *Drosophila melanogaster*

The publicly available datasets used are listed in Table 2. For bulk tissue RNA-seq datasets, the author-processed datasets (expression matrix and differentially expressed gene analysis by either EdgeR or DESeq) were directly used for customized analyses and visualization in this study. For single-cell RNA-seq datasets, the author-annotated expression matrices were directly used for similarity analysis (see details below).

**Table 2.** References and identifiers of the sequencing datasets used in this study.

| **Dataset** | **Reference** | **Identifier** |
| --- | --- | --- |
| Antennal WT bulk RNA-seq of *Drosophila melanogaster* | (Barish et al., 2018) | GSE75986 |
| PN WT bulk RNA-seq of *Drosophila melanogaster* | (Li et al., 2020b) | GSE140093 |
| ORN single-cell RNA-seq of *Drosophila melanogaster* | (Li et al., 2020a; McLaughlin et al., 2021) | GSE162121<br>GSE143038 |
| PN single-cell RNA-seq of *Drosophila melanogaster* | (Li et al., 2017a; Xie et al., 2021) | GSE161228<br>GSE100058 |
| Amos mutant vs WT antennal RNA-seq of *Drosophila melanogaster* | (Mohapatra and Menuz, 2019) | PRJNA532415 |
| *Atonal* mutant vs WT antennal RNA-seq of *Drosophila melanogaster* | (Menuz et al., 2014) | N.A. |
| *Or* mutant vs WT antennal RNA-seq of *Drosophila melanogaster* | (Deanhardt et al., 2023) | GSE179213 |
| ORN single-cell RNA-seq of *Aedes aegypti* | (Herre et al., 2022) | GSE192978 |
| ORN single-cell RNA-seq of *Ooceraea biroi* | (Brahma et al., 2023) | SRX21504666, SRX21504667 |

### Pairwise similarity analysis for gene combinatorial expression in ORNs and PNs

The ORN single-cell RNA-seq dataset was first filtered to keep cells annotated to a single ORN class. Then, for cells in each stage, we extracted their *beat/side* expression profile, i.e., the vector of which each element is the expression value of each *beat/side* gene. We further filtered out the cells with zero expression for all *beat/side* genes. The remaining cells were used to calculate the expression correlation for *beat/side* genes and other gene combinations, e.g., IgSF-encoding genes or CSM-encoding genes. We first calculated the Spearman’s correlation matrix for *beat/side* expression across the selected cells in each stage (Lv et al., 2022). Then, we assigned the kinship for each cell-cell pair based on the annotated ORN/glomerular identity, sensillum where they are housed, and the sensillar type (Couto et al., 2005; Fishilevich and Vosshall, 2005; Grabe et al., 2016; Marin et al., 2020; McLaughlin et al., 2021; Silbering et al., 2011; Task et al., 2022). Specifically, if two cells are annotated as the same ORN class, this pair is defined as “within ORN class”; if two cells are annotated as different ORN classes but both in the same sensillar subtype, this pair is defined as “within sensillar subtype”; if two cells are annotated as different ORN classes from different sensillar subtypes, but belong to the common sensillar type, this pair is defined as “within sensillar type”; if two cells are annotated as different ORN classes from different sensillar types, the pair is defined as “between sensillar types”.

The PN single-cell dataset was first filtered to keep cells annotated to a single PN class (excluding a small portion of anterior paired lateral (APL) neurons) and corresponding lineage identity. We only kept cells annotated to either adPN or lPN lineage (removed multiglomerular vPNs). Then, we extracted their *beat/side* expression vector for cells in each stage. We further filtered out the cells with zero expression for all *beat/side* genes. The remaining cells were used to analyze different gene combinations at the same stage. We first computed the Spearman’s correlation matrix of *beat/side* expression across the selected cells in each stage. Then, we assigned the kinship for each cell-cell pair based on the annotated PN/glomerular identity and lineage identity. Specifically, if two cells are annotated as the same PN class, this pair is defined as “same glomerulus, same lineage”; if two cells are annotated as different PN classes but in the same PN lineage, this pair is defined as “different glomeruli, same lineage”; if two cells are annotated as different PN classes from different PN lineages, this pair is defined as “different glomeruli, different lineages”. Cell-cell pairs belonging to “same glomerulus, different lineages” were scarce and removed before statistical tests.

We finally plotted the distribution of the correlation coefficient of each kinship category and compared the mean between the two categories using the Mann–Whitney U test. This analysis was expanded to the correlation of IgSF and CSM genes, randomly selected genes, or the whole transcriptome (all genes) at each stage. As the numbers of pairs (n) in each kinship category vary dramatically, we also randomly selected the same number of pairs in each kinship category. Then, we performed statistical tests on the resampled data and found the results to be true. Therefore, all related figures in this paper were plotted based on the original data without resampling.

Shuffled control panels were calculated under the same pipeline as the abovementioned, except the annotations of each cell, ORN/PN class name, and corresponding lineage information were shuffled. We shuffled 1000 times with different random seeds and plotted the average mean of each shuffling run.

A similar analysis was also applied to the *beat/side* genetic labeling results, where the binary expression matrices (with only 0 and 1) were used as the input. In this case, as each glomerulus represents the whole ORN/PN classes, we would not have the “within ORN class” or “same glomerulus, same lineage” category.

### Ortholog identification and phylogenetic analysis of *beats* and *sides* in yellow fever mosquitoes and clonal raider ants

To identify the orthologs of *Drosophila melanogaster beat/side* genes in *Aedes aegypti* (yellow fever mosquitoes) and *Ooceraea biroi* (clonal raider ants), we used two ways to cross-validate the candidate list. For the biased way, we first queried the *Drosophila* genes one by one to get the putative orthologs in the other two species on https://www.orthodb.org (Tegenfeldt et al., 2025). Then, we queried the mosquito and ant genes to get the fruit fly orthologs. This reciprocal approach gave rise to a primary list of candidate *beats/sides* in mosquitoes and ants. For an unbiased way, we used the blastp tool on NCBI (https://blast.ncbi.nlm.nih.gov/Blast.cgi?PAGE=Proteins) to find the orthologous sequences encoded by the mosquito and ant genomes of *Drosophila* Beat-Ia and Side proteins. We applied additional filters, including protein size and AlphaFold-predicted structures (https://alphafold.ebi.ac.uk). We only kept candidates with comparable amino acid lengths to *Drosophila* homologs and similar protein structures (two Ig domains for Beats; five Ig domains and one fibronectin domain for Sides). This gave rise to a longer list of candidates. We compared these two lists and found that the first is contained in the second one. We then ran multiple sequence alignments of all Beat candidate and Side candidate proteins, together with the fruit fly ones, to build the phylogenetic trees, respectively, through the Clustal Omega program (Madeira et al., 2024) using the default parameters on https://www.uniprot.org.

### Single-cell RNA-seq analysis of *beat/side* ortholog expression in the ORNs of yellow fever mosquitoes and clonal raider ants

Author-annotated single-cell RNA-seq datasets of *Aedes aegypti* and *Ooceraea biro* from two prior studies (Table 2) were directly used for downstream analyses and visualization. We extracted the mosquito and ant *beat/sides* genes from the original expression matrices. Based on the phylogenetic reconstruction described earlier, we renamed them to reflect their phylogenetic relationships with analogous genes in fruit flies (Table S1).

### Statistics and code availability

R 4.3.1 and the necessary packages were used for all the abovementioned analyses and visualization. Statistical tests were performed with the built-in functions of R or through GraphPad Prism 9 software. The customized codes to replicate all illustrations and statistical graphs of this study will be uploaded to https://github.com/volkanlab.

---

**Table S1.**
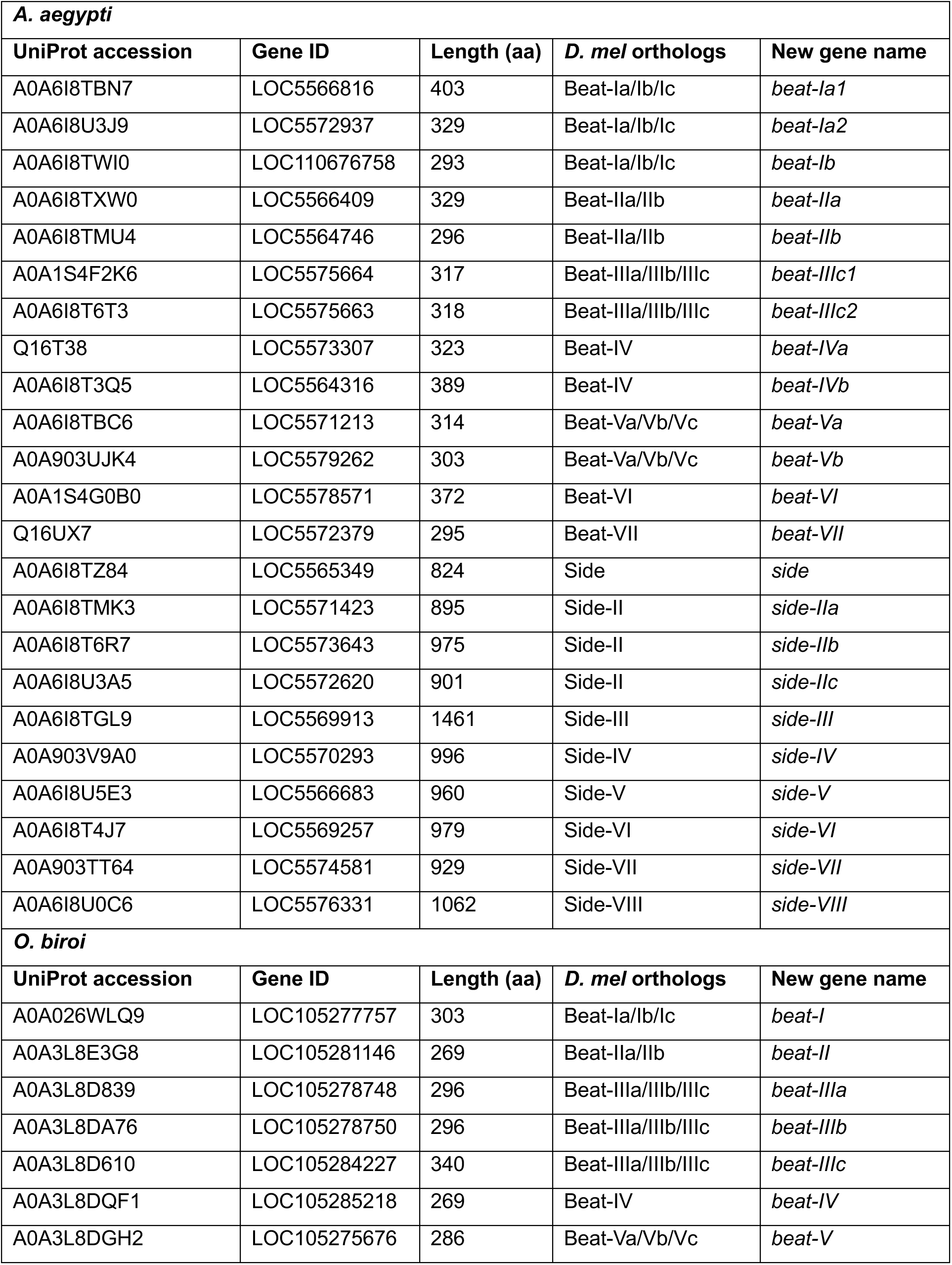
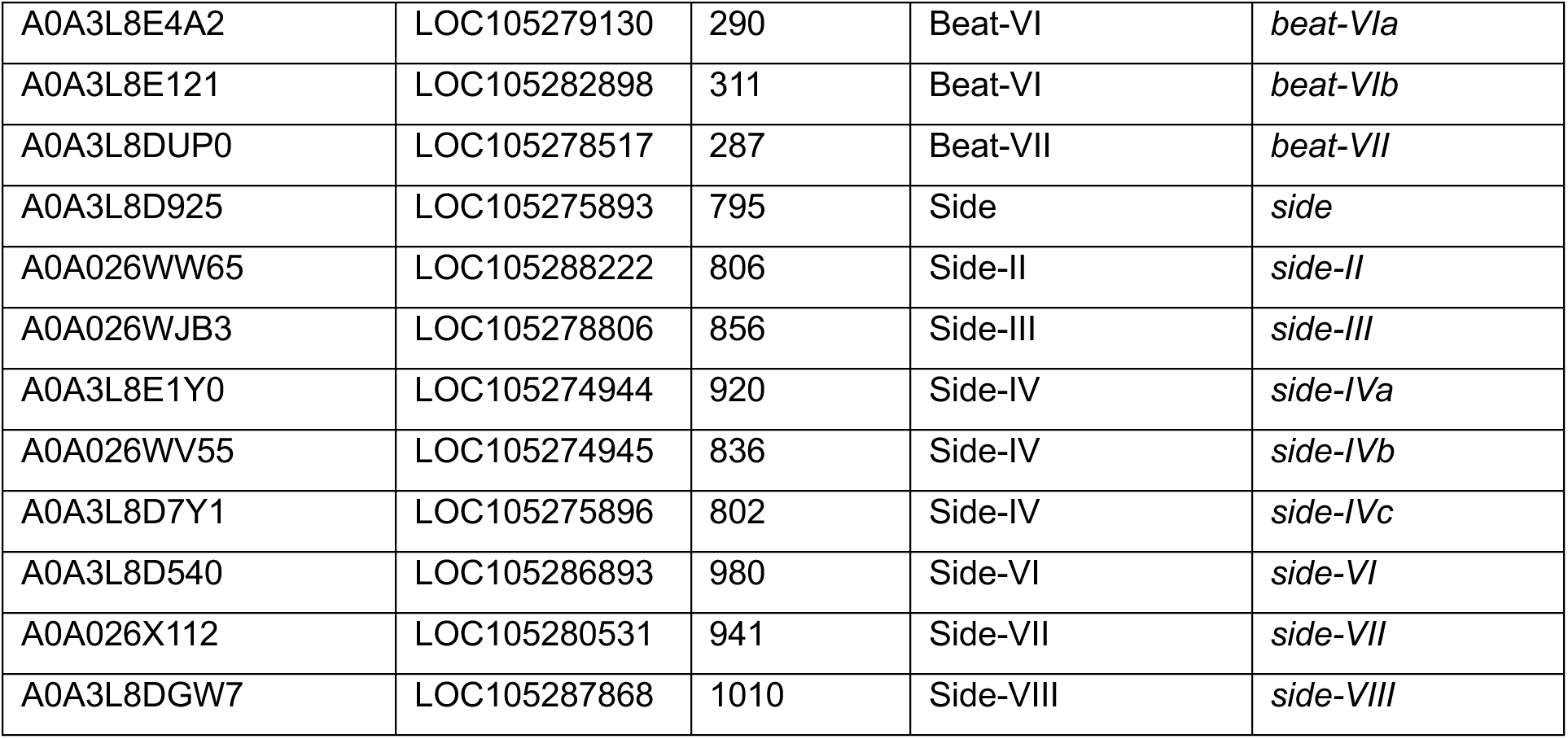
Orthologs of *beats/sides* in yellow fever mosquitoes *Aedes aegypti*, and clonal raider ants *Ooceraea biroi*.

## Author contributions

QD and PCV conceptualized the project and designed the experiments. QD did experiments, analyzed data, and prepared figures. Y-CDC, SO, QD, & LQR generated reagents. SO, RE, CY, and KMV helped with experiments. QD and PCV wrote and edited the manuscript.

## Acknowledgement

The authors would also like to thank the Bloomington Drosophila Stock Center and the Vienna Drosophila Resource Center for providing the fruit fly strains and the Duke Light Microscopy Core Facility for technical support for imaging. The authors would also like to thank Drs. Claude Desplan and Richard S. Mann for sharing reagents, and members of the Volkan lab for discussions and comments on the manuscript. The authors particularly thank Allison Carson and Nick Manfred for assistance with the experiments. The authors also thank Drs. Hiroaki Matsunami, Corbin Jones, David McClay, and Amy Bejsovec for stimulating suggestions.

